# The Optic cup is actively shape programmed by independently patterned apical forces

**DOI:** 10.1101/2025.08.21.671431

**Authors:** Ana Patricia Ramos, Lior Moneta, Alicja Szalapak, Louise Dagher, Malcolm Hillebrand, Carl D. Modes, Caren Norden

## Abstract

During morphogenesis, initially flat tissues often must transition into complex 3D shapes, reminiscent of shape-programmable systems in physics and engineering. One key question in developmental biology and physics alike is therefore how distinct mechanical inputs interact to produce the correct shape. To investigate this, we here study the onset of optic cup invagination during vertebrate eye development, combining 3D shape analysis, perturbation experiments, and physical modelling inspired by shape-programmable materials. We find that basal invagination is initiated at the apical surfaces through active, patterned cell behaviours. These behaviours generate in-plane strain patterns reshaping the apical surface and thereby bending the tissue basally. Surprisingly, this means that basal shape initiation is driven by apical dynamics. While retinal pigmented epithelium and neuroepithelium exhibit distinct shape transitions, these are temporally coordinated to jointly shape the optic cup. These findings highlight that 3D shape can emerge from local apical activity of two coordinated patterns.

## Introduction

Shape acquisition is the process by which biological tissues or synthetic materials progressively transform into a target 3D structure. Since shape is often linked to the function of a developing organ or of an engineered functional material, understanding shape acquisition in 3D is a long-standing challenge in biology and physics alike. In recent years, it has become increasingly clear that many of the principles of shape formation in one of the disciplines are shared with the other. Hence, the realm of complex shape acquisition offers a conceptual framework for mutual insights and inspiration between biology and physics.

Synthetic materials can be transformed from simple 2D or quasi-2D form into a complex 3D target shape using a variety of strategies^1^. One strategy to induce controlled shape transitions involves active shape-changing materials – such as hydrogels or liquid crystal solids – that rely on material heterogeneity or anisotropy to achieve out-of-plane curvatures driven by non-uniform growth in response to external stimuli^2–4^. Here, shape programming becomes possible through the “encoding” of the target shape into a 2D sheet, allowing for precise control over when and how the material re-shapes itself.

During embryonic development, shape emergence during tissue morphogenesis and organ formation is orchestrated by the tightly coordinated interplay between physical, cellular and molecular mechanisms. To explain how this interplay can generate form, 3D shape transitions are often categorized into two main classes: 1) instabilities mediated by external forces that can be transmitted either through neighbouring tissues or the extracellular matrix. This is seen, for example, during the formation of the cephalic furrow or wing disc doming in drosophila^5,6^; 2) apical-basal changes mediated by local differences in mechanics. This is seen, for example, during vertebrate neural tube closing or sea urchin gastrulation^7,8^. While these two classes of shape emergence have helped to elucidate the formation of diverse structures, they are insufficient to explain the full array of 3D morphogenetic events. Consequently, recent studies have uncovered an additional class of morphogenetic mechanisms that build on the concepts from shape-programmed materials mentioned above^9–11^. Here, patterned in-plane spontaneous strains can drive specific shape outcomes. In this context, active collective cell behaviours, such as oriented proliferation^9^ or cell rearrangements^10^ (also known as neighbour exchanges or t1 events), provide such spontaneous strains by changing the preferred lengths within the tissue. These internal, tissue-intrinsic length changes can, in turn, alter tissue curvature, a fundamental component of shape change.

With these different classes described for shape formation in single epithelia, it becomes more feasible to probe how more complicated progressively shaping tissues respond to or incorporate the presence of multiple simultaneous morphogenetic inputs. Indeed, in complex structures with several layers and/or cell types, cell and tissue rearrangements often occur in parallel, which can lead to multiple cooperative or competitive mechanical couplings among different tissue elements and/or deformation modes. These couplings must presumably be tightly coordinated to achieve a target shape in a robust and reproducible manner. While this complexity is present in many developmental contexts and across shape-programmable systems more generally, it is still not understood how multiple mechanical mechanisms interact to deliver robust target shapes.

To gain further insight into the emergence of shape in such complex developmental frameworks, we here investigate the formation of the Optic Cup (OC), the precursor of the vertebrate eye. We apply a combination of theory and modelling from the physics of shape-programmable systems with developmental biology. The developing zebrafish is used as a model, as here the behaviour of cells and tissues can be captured in their native 4D conditions. During OC formation, the flat Optic Vesicle (OV), consisting of two apposing epithelial layers, invaginates to shape the future eye^12,13^ (Supplementary Video 1). The apical surfaces of these layers face inwards and are in close proximity to each other (Fig. 1a). Invagination occurs at the basal surface of the distal layer, which will give rise to the Neuroepithelium (NE). While the tissue invaginates, several morphogenetic events occur simultaneously (Fig. 1a): in the proximal layer, cells of the prospective Retinal Pigmented Epithelium (RPE) flatten^14–17^; at the opposite side, the invaginating distal layer NE cells undergo basal constriction^18–20^; and rim cells at the margins actively migrate from the RPE into the NE^14,20–22^. At the same time, the lens placode, a separate ectodermal tissue adjacent to and facing the OV, begins to move and protrude into the forming OC^23,24^.

**Fig. 1.**
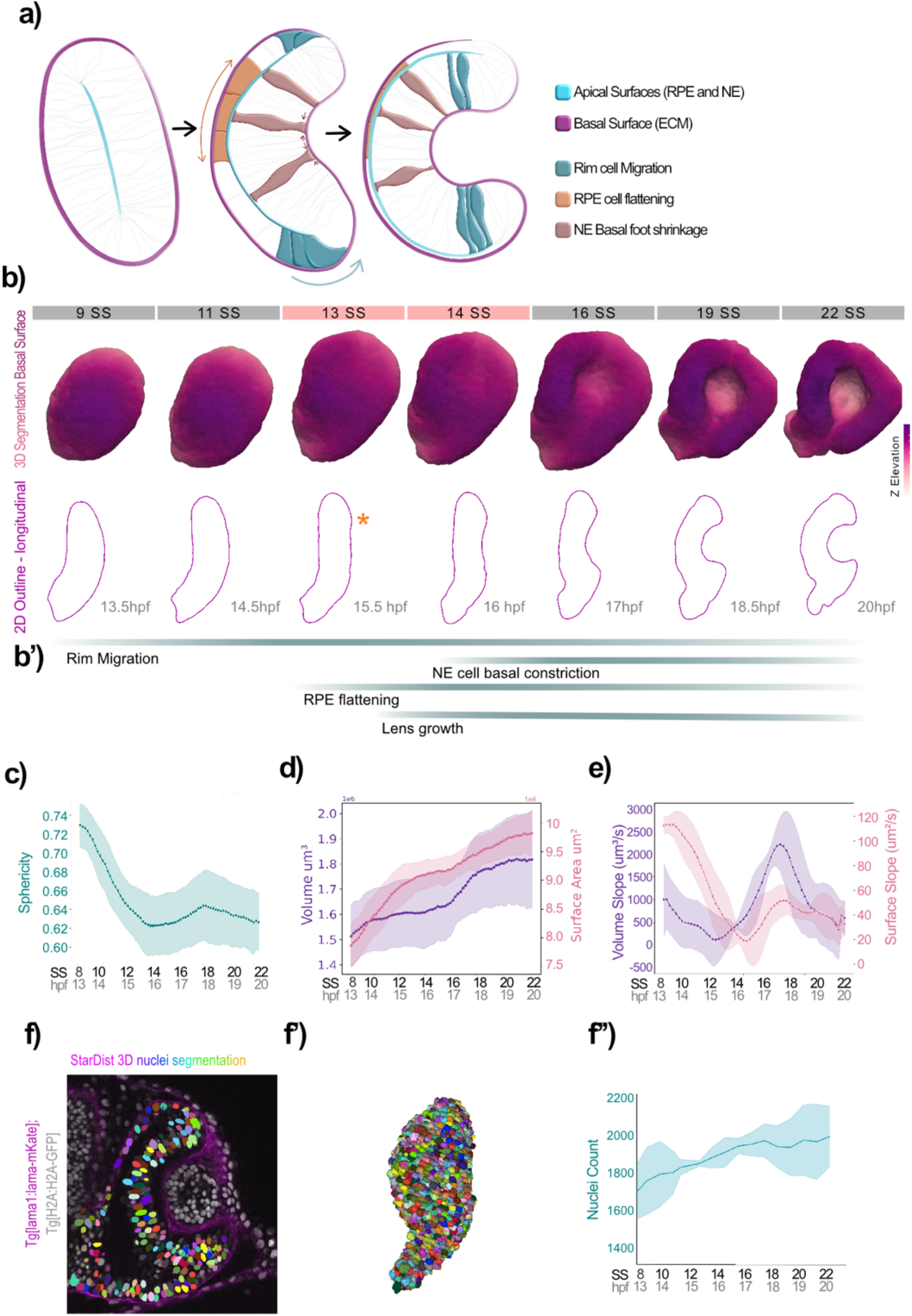
Shape changes during OC formation. **a**, schematic representation of the rearrangements occurring throughout Optic Cup Formation: rim cell migration, Retinal Pigmented Epithelium (RPE) cell flattening, Neuroepithelium (NE) basal shrinkage. **b**, basal surface segmentation in 3D over Optic Cup Formation, and respective 2D longitudinal sections. Developmental time is indicated in somite stage (SS) and hours post-fertilization (hpf). The onset of invagination is marked by the asterisk **b’** time frame of the cell rearrangements depicted in (**a**), in relation to the 3D optic cup shape changes. **c**,**d**,**e**, geometrical measurements obtained from the 3D segmentations, over time; dark lines represent the mean values and lighter regions depict the standard deviation (n=3 embryos), developmental time is indicated in somite stage (SS) and hours post-fertilization (hpf); **c**, mean sphericity of the OC (calculated as ration of volume to surface area (V/A^^(3/2)^); **d**, total volume in um^3^ (dark purple) and surface area in um^2^ (magenta); **e**, mean slope of the volume (in um^3^/s) and of the surface area (in um^2^/s), sliding time window of 15 timepoints. **f-f’’**, nuclear segmentation and quantification in the OC, using Stardist; **f**, overlay of optical slice through the Optic Cup. Nuclei are labelled with H2B-GFP (in grey), extracellular matrix (ECM) is labelled with laminin-mKate (magenta), segmented nuclei are represented by multi-coloured labels. Segmentation of the ECM was used to exclude the segmented nuclei outside of the optic cup; **f’**, 3D reconstruction of the segmented nuclei depicted in (**f**); **f’’**, quantification of the number of nuclei within the forming optic cup dark line represents the mean value, lighter regions depict the standard deviation (n=3 embryos). Developmental time is indicated in somite stage (SS) and hours post-fertilization (hpf).

Several hypotheses have been put forward to explain OC formation *in vivo* and in *ex vivo* developing retinal organoids. In these, invagination is either achieved through an instability mediated by external forces, such as buckling instabilities due to ECM constraints^25,26^, or through basal to apical area changes, namely basal foot constriction^18^ and apical expansion^27^. However, so far studies have mostly focused on separate, individual events, without exploring the potential interplay between them. Whether this interplay is needed to drive the *onset* of OC invagination remains thus an open question, as the exact mechanism of this shape transition is still not fully understood

We therefore here focus on this *onset* of OC invagination to uncover which tissue rearrangements contribute to it and how they do so in a spatiotemporally resolved manner. In order to reach a comprehensive overview of the parallel processes that contribute to OC formation, we consider how 3D shape transitions in the different epithelia are associated with the overall shape of the forming OC. By integrating quantitative 3D shape measurements with theoretical modelling, we show that OC invagination is initiated at the apical surfaces at the interior of the tissue. We find that active in-plane collective cell behaviours at the apical surface of both the RPE and NE layers generate the patterned spontaneous strains that lead to apical curvature. This apical curvature onset is then sufficient to drive NE basal surface invagination. We further demonstrate that the apical surface of both layers (RPE and NE) act independently of each other to initiate their own shape transition while together they contribute to the final OC shape.

Overall, our work not only offers a new perspective on OC formation but also promotes a broader understanding of morphogenesis by identifying how independent patterns of morphogenetic activity work together to give rise to a 3D organ. In addition, this work advances the biophysics of shape mechanics by considering interacting shaping mechanisms and shedding further light on a newly identified modality of in-plane shape programming driving biological shape transitions.

## Results

### Quantitative analysis to extract shape changes during optic cup formation

As the Extracellular Matrix (ECM) surrounds the entire OC we used it as a proxy to quantify shape changes during OC formation^21^ (Extended Data Fig. 1). We made use of the double transgenic tg(lama2:lama-mKate; RX2:GFP-caax) line, in which the ECM protein Laminin and the cell membranes of the OC are labelled. Developing embryos were imaged between 8 to 20 somite stage (SS), the time window that captures the full extent of shape changes during the transition from OV into OC.

The laminin signal was segmented over time, and a triangulated mesh was created from this segmentation (Fig. 1b and Supplementary Video 2) (details in Extended Data Fig. 3a and methods).

Analysis of the curvature progression in these 3D segmentations, showed that OC invagination starts between 13 and 14 SS (Fig. 1b). This OC invagination is preceded by a shape inversion, as the OV is initially convex at 8 SS. In line with this observation, the sphericity of the OV gradually decreases from 8 SS until the onset of invagination at 13-14 SS (Fig. 1c). This means that the OV starts changing its shape at least two hours before the onset of NE basal invagination.

During the transition from OV to OC, volume and surface area of the emerging cup increase from 8 SS (Fig. 1d,e). Even though growth rates are variable over time, there is a sharp increase of volume and surface area growth rate from the moment the OV starts to invaginate. Quantification of 3D segmented nuclei using a pre-trained Stardist model^28^ (details in methods and Extended Data Fig. 2b) showed that OC volume growth is accompanied by an increase in cell number (Fig. 1f).

Overall, our analysis based on the segmentation of the ECM and nuclei within the OV allowed us to reproducibly quantify geometrical measures and 3D shape changes over OC formation. Our observation that the ECM closely follows the shape changes of the overall tissue, suggested that the ECM does not provide a direct constraint against which the tissue buckles. Together, this initial analysis generated the basis for following 3D quantification of shape changes.

### Retinal Pigmented Epithelium and Neuroepithelium show coordinated curvature onset

Multiple epithelial and cellular rearrangements take place within the developing OC, as well as in the surrounding tissues, while the tissue acquires its shape (Fig. 1a and b’).

To understand which epithelial changes are responsible for OV invagination onset, we related the time sequence of rim migration, RPE cell flattening, NE basal shrinkage and lens invasion to the ongoing shape changes. Basal surface activity in the neuroepithelium (NE), specifically basal cell shrinkage driven by localized actomyosin enrichment, has been suggested to be involved in invagination^18,20^. However, so far NE basal constriction and acto-myosin activity have been studied only after invagination onset. Therefore, it is not yet clear whether this event is sufficient to trigger invagination onset.

To investigate this possibility, we monitored Actin and Myosin distribution during the OV to OC transition. Protein distribution was quantified by fluorescence intensity analysis of GFP-tagged Myosin and Utrophin, or of the GFP-tagged Actin chromobody (details in methods and Extended Data Fig. 2c). We found that the enrichment of Actin and Myosin at the NE basal surface is seen from at least 10 SS. This correlated to around two hours before OV invagination onset (Extended Data Fig. 2a,b). In addition, local foci of Actin do not correlate with the invaginating region (Extended Data Fig. 2c). Therefore, while basal acto-myosin activity could play a role in invagination progression, our observations that the acto-myosin basal bias is present before invagination onset and does not change over time make it unlikely to serve as an initiator of OV invagination.

The lens placode starts to grow from the adjacent ectoderm at the same time as the NE basal surface starts to invaginate, making it a further candidate to be involved in the onset of OV invagination by pushing the NE inward^23,29^. To test this hypothesis, we used an UV laser and induced cuts in the lens placode region, perpendicular to the potential pushing direction, at 15-16SS. When monitoring the NE basal surface curvature, we observed that, upon ablation, the inward curvature of the OC increased. This is the opposite effect of what would be expected if the growing placode was pushing the NE inwards (Fig. 2a and Supplementary Video 3, n=5 embryos). This finding implies that the growing lens is not actively pushing the NE inwards. Instead, our data suggests that the NE could be pulling the lens into its eventual position at the centre of the invaginating NE. We thus concluded that also pushing forces by the incoming lens are not likely to be the initial driver of NE invagination onset.

**Fig. 2.**
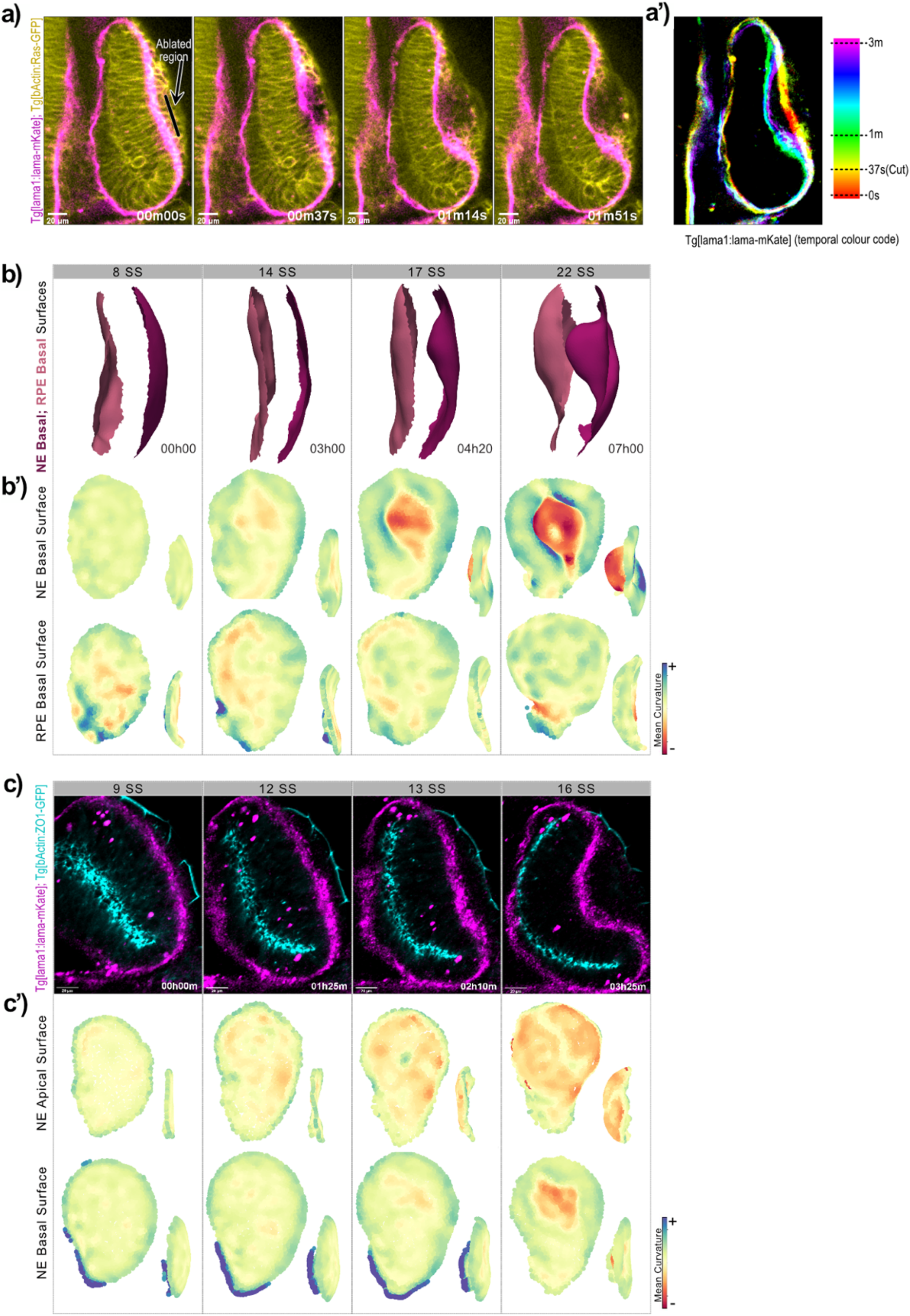
OC invagination starts at the inner apical surfaces. **a**,**a’**, lens placode cells ablation, spinning disk optical slice. ECM is labelled with lama-mKate (magenta) and cell membranes with ras-GFP (yellow). Black line depicts the site of the ablation (before ablation). Images were taken every 37 sec. **a’**, colour coded projection of the ECM over time, from before ablation (red) until 3 minutes after ablation (purple) **b**,**b’**, Retinal Pigmented Epithelium (RPE) and Neuroepithelium (NE) basal surface curvature progression; **b**, 3D segmented basal surfaces of the OC. The rim regions were removed to be able to analyse the RPE (light pink) and the NE (magenta) separately; **b’** mean curvature quantifications of the mesh points in **(b)** over time, frontal view and side view (inlet). Developmental time is indicated in somite stage (SS) **c, c’** OC basal vs apical surface curvature progression; **c**, confocal optical slice of the developing OC. Basal surface (ECM) is labelled with lama-mKate (magenta), and apical surface (tight-junctions) are labelled with ZO1-GFP (cyan) **c’**, Mesh points mean curvature quantification over time of the apical surface and of the NE basal surface, frontal view and side view (inlet). Meshes were obtained from the segmentation of the basal surface and of the apical surface. Developmental time is indicated in somite stage (SS)

Another tissue event that coincides with OV to OC shape changes is the flattening of the RPE cells. RPE flattening starts between 10 and 14 SS^17^ (Fig. 1b’). While it has been proposed that this process participates in OV invagination^16,17^, its specific contribution to this shape transition is not yet clear. To understand if RPE flattening could be linked to the onset of OV invagination, we compared the progression of shape changes occurring at the basal surface of the NE and RPE. To this end, we separated the segmented basal surface of the RPE from the segmented NE basal surface (Fig. 2b and Supplementary Video 4). This was achieved by identifying and removing the vertex points belonging to the rim region, which disconnected the segmentation of the two surfaces (details in methods and Extended Data Fig. 2d). By quantifying curvature onset and progression for each surface separately, we found that the RPE layer closely follows the curvature patterns of the NE: Convex at 9SS and becoming concave at 14SS, after shape inversion as seen for the NE (Fig. 2b’). This shows that RPE and NE basal surface shape changes occur concurrently. However, the spatial distribution of curvature differs between the two tissues: while the RPE mean curvature is globally uniform, the NE mean curvature is more localised, showing a negative curvature peak at the centre, surrounded by regions of positive curvature (Fig. 2b’). Due to the observed synchronisation in basal surface curvature progression between the RPE and NE it we considered it unlikely that one surface is driving shape changes of the other.

Combined, our results show that neither acto-myosin activity at the NE basal surface nor lens growth are major initiators of OV invagination. The concurrent curvature progression between the RPE and NE basal surfaces further makes it unlikely that activity at the RPE basal surface is driving shape changes at the NE.

### Optic Vesicle invagination starts at the apical surface

In zebrafish, the apical surfaces of the RPE and NE are in close contact to each other. Given the concurrent shape progression between the RPE and NE (Fig. 2b), we speculated that the two close apical surfaces could be driving or coordinating basal surface shape transitions. If this was the case, the apical RPE and NE surfaces should curve before the onset of basal NE invagination. To quantify curvature changes at the basal vs apical surfaces, we followed the ECM and the apical tight junctions, by imaging double transgenic embryos tg(lama:lama2-mKate, tg(bactin:ZO1-GFP) (Fig. 2c and Supplementary Video 5). ECM signal was segmented as described above. Given that RPE and NE apical surfaces remain in close proximity, and are indistinguishable in our images, we segmented the ZO1 signal as a single surface (details in Methods). We then quantified the mean curvature over time at the NE basal surface and at the apical surface. Analysis of curvature progression revealed that, indeed, the apical surface starts to curve 30 to 45 min before the basal NE starts to invaginate (Fig. 2c’ and Supplementary Video 5).

The finding that the apical surfaces start to curve before the basal NE surface, suggests that apical surface activity could be driving the onset of basal invagination.

### Theoretical model of active apical surface as a shape programmed sheet

To start testing our hypothesis that the apical surface is driving basal surface invagination, we first set out to understand how the apical surfaces themselves could be driven into their target shapes. For this, we used TopoSPAM^30,31^, a simulation framework that allows the modelling of possible physical mechanisms of shape-programmed transitions (Fig. 3a). The resulting simulations were then used to inform experiments, and vice-versa, creating a feedback loop between experiment and simulation scenarios.

**Fig. 3.**
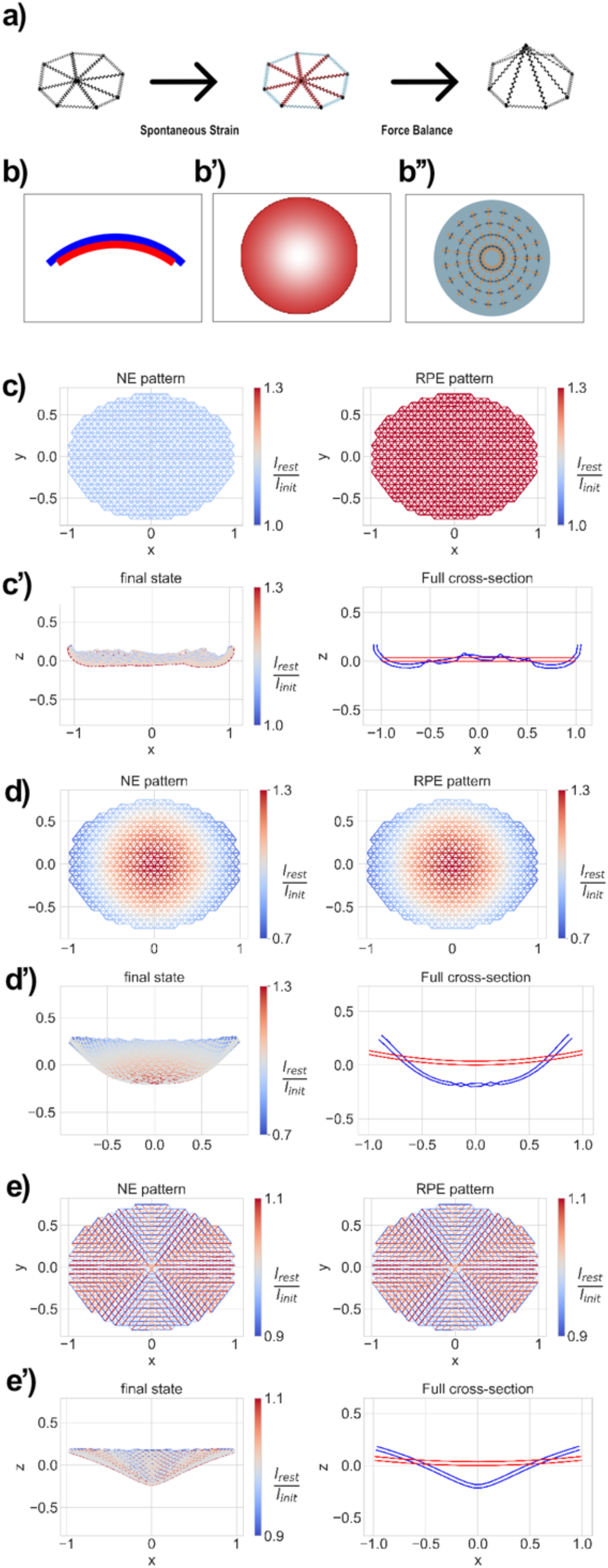
Simulations of the theoretical candidate patterns for RPE and NE apical surface cupping. **a**, The optic cup is modelled as an elastic spring network. The network is initialized in a stress-free state. We apply a spontaneous strain field that changes the springs’ rest length, which is then relaxed to achieve a new 3D configuration of the network. **b-b’’**, Illustration of three cupping mechanisms **b**, total area mismatch between the layers, the red layer has a larger total surface than the blue which leads to cup formation through geometric instability; **b’**, in-layer local isotropic growth gradient - here cells closer to the centre of the layer have a higher increase in area than cells closer to the edge, leading again to a geometric instability that forms a cup; **b’’**, an in-layer uniform local anisotropic strain, where cells grow radially but shrink tangentially resulting in an instability that leads to cup formation. **c-e**, spontaneous strain pattern candidates at the Neuroepithelium (NE) and Retinal pigmented epithelium (RPE), and respective simulation results on the simplified elliptical meshes. All three lead to an invagination of the initial state; **c**, strain pattern resulting from a total increase in cell area at both layers, as depicted in **(b); c’**, final 3D configuration of the mesh after relaxation (full 3D and cross section); **d**, strain pattern resulting from an in-plane isotropic gradient of cell area increase at both layers, as depicted in **(b’); d’** final 3D configuration of the mesh after relaxation; **e**, strain pattern resulting from an in-plane local anisotropic strain, as depicted in **(b’’); e’**, final 3D configuration of the mesh after relaxation.

To model the potential physical mechanisms by which the apical surfaces could acquire their shape during OC formation, we created a boundary-free triangular network of springs, with a simplified ellipsoid geometry. Given that the apical surfaces of the OV are in close contact with each other and show no lumen in between them, our network includes two mechanically connected layers. By then applying different spontaneous strain patterns to the initially stress-free networks, TopoSPAM updates each spring rest length, accordingly, leading to a shape change upon relaxation of the imposed stresses (details in Methods).

While several mechanical mechanisms could lead to curvature emergence on a flat sheet, we focused on the response of our network to three candidate spontaneous strain patterns, that are most likely to apply in our biological context: 1) Total area mismatch, an apical-apical analogue to the more traditional apical-basal contractions; 2) Non-uniform area growth, as could occur from non-uniform tissue thinning and 3) Radial anisotropic strain, as could be produced, for example, by cell neighbour rearrangements (Fig. 3b).

In our simulations for 1), total area mismatch was achieved by applying a global isotropic growth with different magnitudes on each layer/surface (Fig. 3c). In this scenario, a mismatch between the areas of the two layers leads to a geometric incompatibility. This, in turn, forces the network to undergo a shape transition from a flat to a cup-like configuration. In scenario 2), we imposed a radial gradient of isotropic growth on each surface (Fig. 3d). Here, the differential growth across the surface causes the central region to experience higher growth than the periphery. This results in an incompatibility across the initially flat sheet which in turn leads to internal stresses that drive a shape transition towards a spherical configuration. The same pattern was applied to both layers, which then act independently of each other to reach their respective curvature. To reach anisotropic growth for scenario 3), we applied a constant radial extension and tangential contraction for each layer (Fig. 3e). Considering a circle taken around the centre of the layer, this pattern leads to an increase in the circle’s radius while shortening its circumference. This causes a geometric incompatibility that is, once more, resolved by the formation of a curved cup-like structure.

Together, our simulations showed that, as expected, all these mechanisms are capable of driving shape changes. However, due to the fact that our initial network geometry was a simplified representation of the apical surface shape, we decided to test the effect of the three strain patterns on the actual apical surface geometry of the OV. For this we built an effective network representation of the apical surface, by using as a scaffold the apical surface segmentation. This network with the updated geometry was then used as the initial stress-free condition, to which we applied the above-mentioned strain patterns. In all three scenarios, the network was able to reach cup-like shapes (Extended Data Fig. 4). Thus, our modelling approach confirms that three different physical mechanisms could drive apical surface shape transitions: total area mismatch, non-uniform area growth and radial anisotropic growth.

### Retinal Pigmented Epithelium and Neuroepithelium apical surfaces curve independently

Our network simulations showed three possibilities by which apical surfaces could curve. To test whether any of these simulated mechanisms was indeed involved in driving apical surface curvature we used our experimental frameworks.

In a biological context, shape transitions driven by total area mismatch, as proposed in scenario 1), can be achieved by changes in total cell number or cell area. This, however, requires that the two layers remain connected to each other as only this would allow for a geometric incompatibility to arise. Connections between two apposing apical surfaces are not commonly described during development, but they have been reported in transient epithelial states during zebrafish neurulation^32^ and in the developing Xenopus eye^33^. To test if the apical surfaces of the RPE and NE were indeed connected, we used an UV laser to make a small incision at the edge of the apical surface. Ablation was performed at 8 SS (before curvature onset) and the behaviour of the apical surfaces was monitored by following the signal of the apical surface marker ZO1. Upon ablation, the apical surfaces of RPE and NE separated at the ablation point. Interestingly, this separation propagated beyond the cut region, towards the opposite end of the OV reminiscent to an opening zipper (Fig. 4a Supplementary Video 6). This behaviour suggested that the RPE and the NE apical layers are indeed connected. In our simulations, disruption of the connection between the two apical layers prevented the network from undergoing proper shape transformation (Extended Data Fig. 5). To test if such separation would impair OV invagination, we monitored curvature progression after separation of the RPE and NE apical surfaces. To our surprise, both RPE and NE curvature progressed normally (Fig. 4b and Supplementary Video 7), with only a small delay in invagination onset when compared to the control, non-ablated, eye. Thus, RPE and NE are able to curve independently of each other. The fact that the distance and coupling between the two apical surfaces is not crucial for invagination onset means that we can rule out the first theoretically proposed mechanism in which the apical surfaces are changing shape via a mismatch of total apical area.

**Fig. 4.**
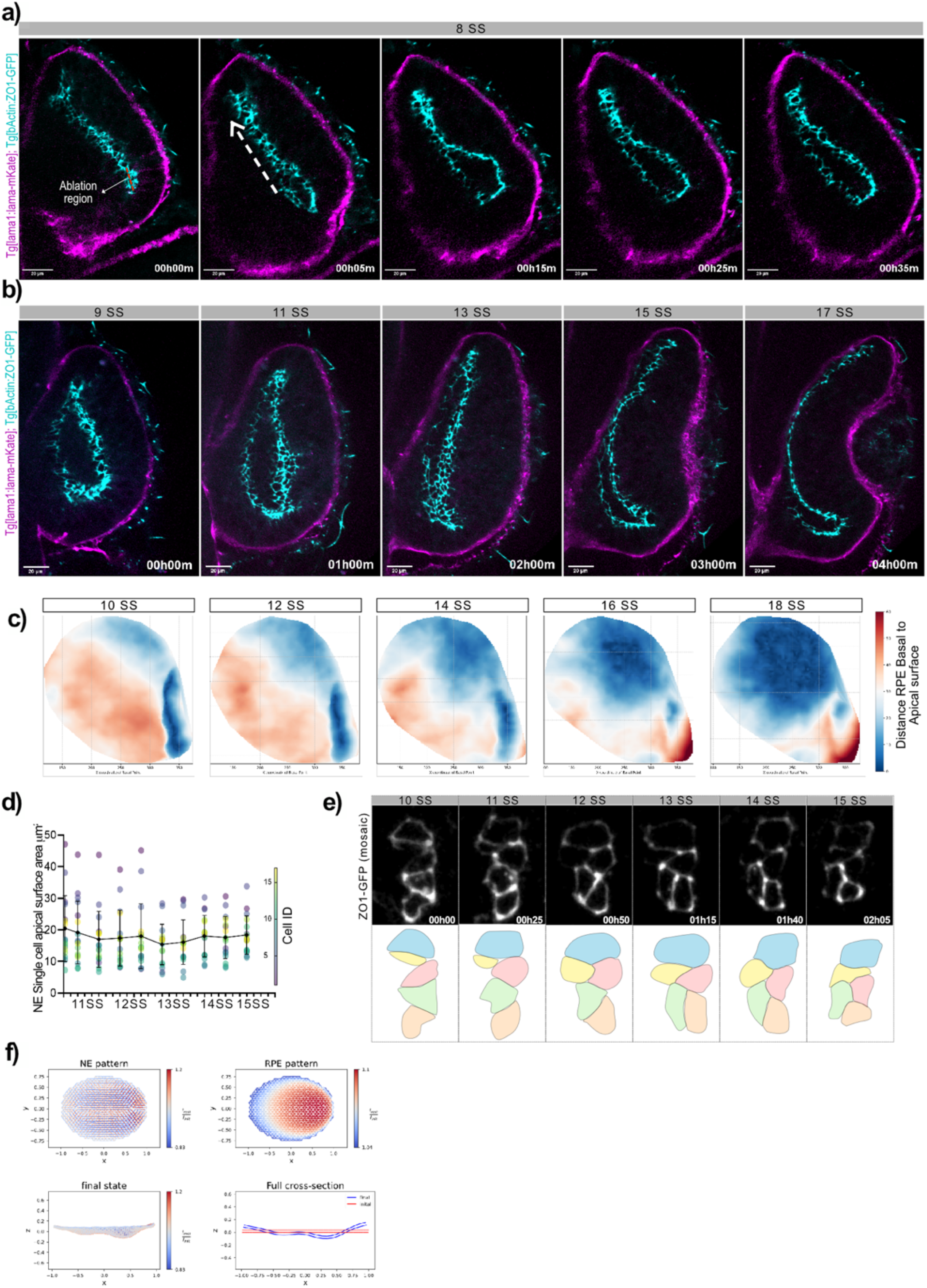
Distinct cell behaviours drive RPE and NE apical surface curvature. **a**,**b**, RPE and NE apical surfaces separation after ablation, spinning disk optical sections over time. Basal surface (ECM) is labelled with lama-mKate (magenta), and RPE and NE apical surfaces (tight-junctions) are labelled with ZO1-GFP (cyan). **a**, Red dashed line depicts the ablated region, white arrow depicts direction of the opening of the two apical surfaces **b**, OC invagination progression after separation of the RPE and NE layers. Developmental time is indicated in somite stage (SS). **c**, distance between Retinal Pigmented Epithelium (RPE) basal surface and apical surface throughout OC Formation. Distance is represented as a color-coded heat map, and was obtained by measuring the shortest distance between 2 points of the RPE basal and apical surface meshes. **d**, Neuroepithelium single cell apical surface area during apical surface curvature onset. Each point depicts the area of a single cell (colour depicts cell ID). The solid line depicts the mean area values, with standard deviation. **e**, ZO1-GFP labelling of a cell mosaic, depicting the apical surface of the cells during apical surface invagination **c’**, schematic of the cells in **(c)**, showing the neighbour rearrangements **f** Realization of our inferred layer specific strain pattern on the apical surfaces in the simplified mesh. A patterned gradient of anisotropic strain on the Neuroepithelium (NE) and a patterned gradient of isotropic growth on the Retinal Pigmented Epithelium (RPE) successfully leads to an invagination of the initial flat state.

### Retinal Pigmented Epithelium and Neuroepithelium use different physical mechanisms to achieve respective apical surface curvature

We next tested scenario 2) of our simulations in which apical surface shape transitions are driven by gradients of isotropic cell area growth. While the OC invaginates, cells in the RPE layer start to flatten and substantially increase their apical surface^17^. This flattening is, however, not uniform through the whole OV, which in turn can lead to a gradient of cell apical surface area increases. To probe the pattern of apical surface area growth, we quantified the progression of cell flattening at the RPE layer. Since cell flattening decreases the distance between the apical and basal surface of the cell, we quantified the distance between the overall RPE basal and apical surface in the respective 3D meshes. This allowed us to build a heat map of apical to basal distances that shows the progression of cell flattening, and the emergence of a gradient from the dorsal-medial region of the OV (Fig. 4c and Supplementary Video 8). The presence of this gradient suggests that indeed RPE apical surface curvature could rely on a gradient of surface area changes.

We then asked if the apical surface area of single NE cells is also changing during apical surface curvature. To answer this, we measured the cell apical surfaces of single NE cells. Small groups of cells were labelled mosaically by injecting ZO1-GFP mRNA in 64 cell stage embryos to be able to differentiate between RPE and NE cells. Cell mosaics were then followed through live imaging in embryos between 11 and 15 SS, the time window of apical surface curvature initiation. We found that while the apical surface areas are variable when comparing different NE cells, there was no trend for the apical area to increase or decrease over time (Fig. 4d). We therefore excluded the possibility that a gradient of apical surface area acts as a driver of NE apical surface shape changes.

To understand how the NE apical surface reaches its target shape, we focused on the mechanism described in scenario 3) in which shape changes are achieved through local anisotropic strain patterns. Such anisotropic patterns can be achieved by different processes. The tissue could either feature anisotropic changes in cell shape; or feature topological events, for example cell rearrangements such as neighbour exchanges. When we followed the apical surface of NE cell mosaics labelled with ZO1-GFP, we observed that frequent cell rearrangements occur at stages at which apical surface curvature initiates (Fig 4D). Quantification of the orientation of these rearrangements were consistent with a bias to an underlying elliptical pattern (Extended Data Fig. 6).

Thus far, our experimental results and theoretical simulations suggest that apical surface curvature onset could be driven by a pattern of gradients of active area growth in the RPE in combination with a pattern of active anisotropic rearrangements in the NE, thus a combination between scenario 2 and 3. Informed by these experiments, we updated our model of apical surface shape transitions to reflect the two different strain patterns on each surface (Fig. 4f). Here, the surface corresponding to the RPE presents a gradient of area growth, while the surface corresponding to the NE presents an anisotropic spontaneous strain pattern. Given the apparent elliptical symmetry observed in the NE rearrangements, we assumed that the active strain patterns are elliptical. Informed by the shape of the apical surface boundary and the apparent elliptical symmetry observed in the NE rearrangements, we further assumed that the active spontaneous pattern on each side would be elliptical, mediated by a pair of +1/2 defects located at the ellipse foci in the anisotropic case.

Taken together, our data points to the existence of different and independent physical mechanisms driving the shape transitions of RPE and NE apical surfaces.

### A theoretical model of active apical surface drives basal surface invagination

We so far narrowed down the potential physical mechanisms that drive apical surface shape transitions. However, it remained to be tested if such apical activity could indeed be the main driver of basal invagination. To theoretically test the hypothesis that OC formation is driven by the coordination of independently established patterns at the RPE and NE apical surface we updated our model. Instead of only two layers corresponding to the RPE and NE apical surfaces, we added two additional layers corresponding to the RPE and NE basal surfaces. As in the previous apical surface model (Fig. 4f), we assumed different active patterns at the RPE and NE to provide the spontaneous strains that lead to apical surface shape transitions. For the basal surfaces however, we assumed that they only respond passively. We further assumed that the connection within basal surfaces and those connecting the basal to apical surfaces are less stiff than those within, and between, the apical surfaces.

To determine if these coupled patterns on the NE and RPE would be sufficient to drive OC invagination, we explored the parameter space associated with the magnitude of the anisotropic spontaneous strain on the NE side and the magnitude and size scaling of the isotropic spontaneous strains on the RPE side (Extended Data Fig.7). Indeed, we found a broad range of parameters over which these mechanisms successfully drive OC formation (Fig. 5a). In addition, applying the same inferred pattern to a network representation built from the apical and basal surface segmentations, resulted in cup formation similar to the biological system (extended Data Fig.8).

**Fig. 5.**
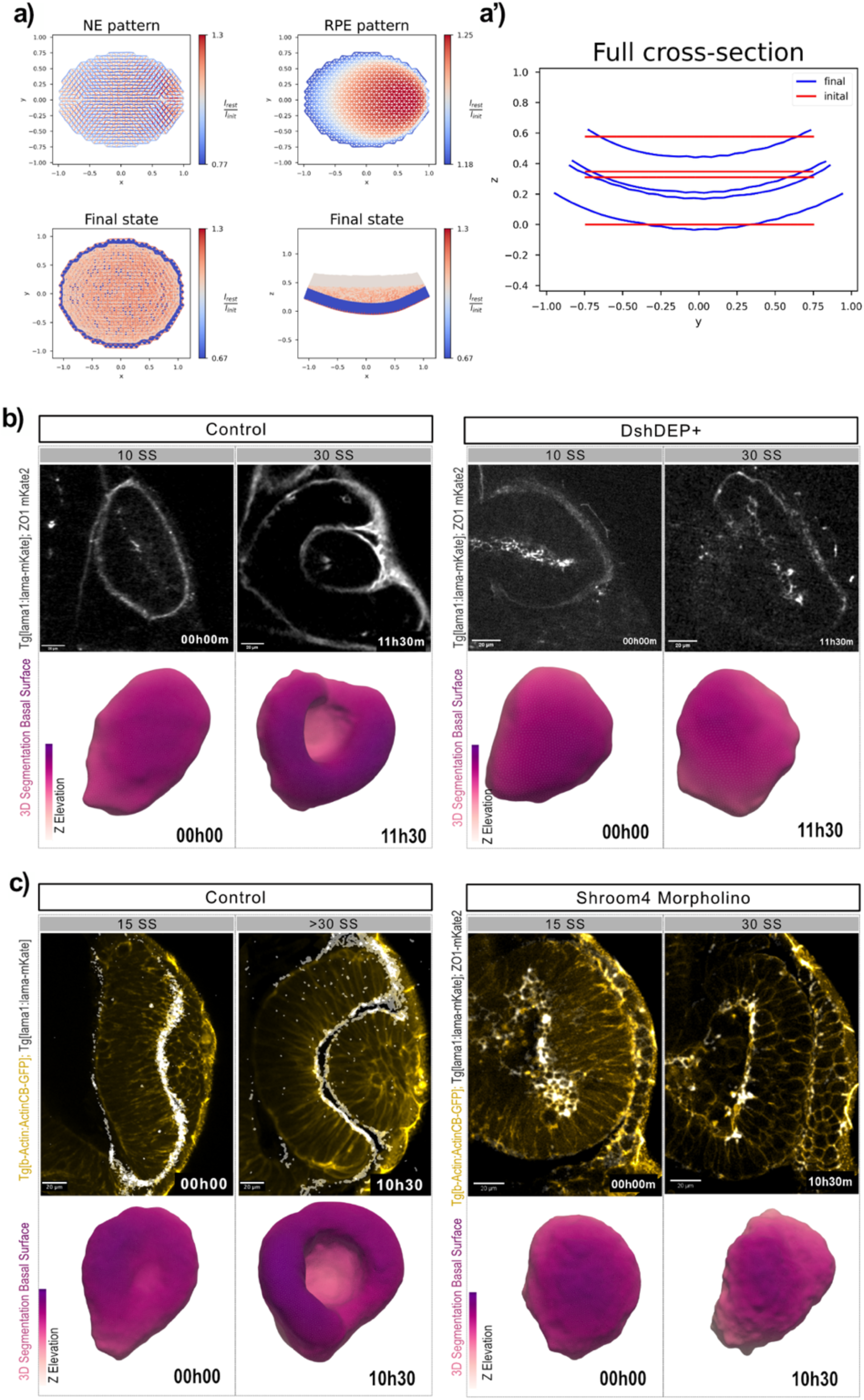
Patterned cell rearrangements at the apical surface drive basal surface invagination onset. **a**, realization of our inferred layer specific strain pattern in the full (four layered) simplified mesh. A patterned gradient of anisotropic strain on the Neuroepithelium (NE) and a patterned gradient of isotropic growth on the Retinal Pigmented Epithelium (RPE) successfully lead to an invagination of the entire network, including the two passive outer layers (corresponding to the NE and RPE basal surfaces).**b**,**b’** disruption of the PCP pathway using dominant negative DSH (DshDEP). **b**, spinning disk optical slices of the forming OC in control and in embryos injected with DshDEP+. Images were taken from a time-lapse movie that lasted 12h. The ECM is labelled with lama-mKate and the apical surface with ZO1-GFP (all grey). **b’** 3D basal surface segmentation of the OC depicted in **(b)**, colour coded for Z elevation. **c**,**c’** disruption of Shroom4 expression **c**, spinning disk optical slices of the forming OC in control and in embryos injected with Shroom4 morpholino. Images were taken from a time-lapse movie that lasted 11h. The ECM is labelled with lama-mKate (grey) and cell membranes are labelled with ActinCB-GFP (yellow). Shroom4 morpholino OCs also show the apical surface labelled with ZO1-GFP (grey); **c’** 3D basal surface segmentation of the OC depicted in **(c)** colour coded for Z elevation.

Together, our experimental data and theory suggest that the RPE and the NE rely on different mechanisms to initiate their own curvature. Coupled together, according to our simulations, this combination would be sufficient to drive the overall formation of the OC.

### Optic vesicle invagination onset is driven by patterned spontaneous strains at the apical surfaces

To experimentally test the idea that NE basal surface curvature can be achieved through the combination of patterned spontaneous strains at the apical surfaces, we aimed to impair one of the spontaneous strain patterns.

As we proposed previously that the cell rearrangements occurring at the apical surface of the NE are the potential source of the in-plane anisotropic strain patterns we expect them to drive apical surface shape changes. To clarify this further we disrupted these rearrangements.

Cell rearrangements including the ones that lead to tissue in plane changes rely on the anisotropic contractility of the cells. Typically, this anisotropy is, at least in part, controlled by components of the planar cell polarity pathway (PCP)^34,35^. Therefore, we aimed to reduce patterned rearrangements in the OC, by disrupting the PCP pathway. For this, the mRNA for a dominant-negative form of the Dishevelled protein (Dsh-DEP+)^36,37^, an important component of the PCP pathway^38,39^, was injected together with ZO1-mKate mRNA in the tg(lama2:lama-mKate) line. OC development was followed through live-imaging from 10SS, for 12 hours. Strikingly, even after 12 hours of imaging, the OV was unable to invaginate at the basal surfaces in 12 out of 16 imaged embryos (Fig. 5b and Supplementary Video 9), while control embryos started to invaginate within two hours of imaging. Importantly, overexpression of Dsh-DEP+ did not lead to overall developmental delays.

To further confirm that the observed phenotype was caused by the disruption of the patterned strains at the apical surface, we interfered with Shroom expression. Shroom is a known downstream effector of the PCP pathway that primarily functions at the apical surface^8,40,41^, controlling the actomyosin machinery. Analysis of previously published RNA sequencing datasets of the developing OC^42^, confirmed that Shroom4, an orthologue of the Shroom family, is expressed during invagination of the OC. Similar to Shroom3, Shroom4 has been previously shown to be an actin-associated protein^43^. Therefore we decided to interfere with Shroom4 expression, by using a previously validated morpholino^44^. Shroom4 MO was injected together with ZO1-GFP mRNA to label the apical surface, into the tg(lama2:lama-GFP) line, at the one cell stage. OC morphogenesis was followed through live imaging for 10 hours. As seen for the experiments using dominant negative dishevelled, also in this condition OV invagination failed to start in 16 out of 18 embryos (Fig. 5c and Supplementary Video 9).

Returning to the model we tested a similar perturbation where we disrupted the anisotropic pattern on the NE apical surface. We found that also here, eliminating spontaneous strains at the NE apical surface impedes OC invagination (Extended Data Fig. 9).

Together these results confirm that the apical surface activity of the zebrafish OV is the major initiator of NE basal surface invagination

## Discussion

The formation of 3D structures from quasi-2D tissues is an important hallmark of shape acquisitions in biology and physics, and a key step for the generation of functional organs and materials. Here, we follow the process of OC formation, a crucial step in eye development that shapes the eye’s curvature, essential for proper light focusing and visual function. Our 3D analysis, which brings together the physics of shape programming and developmental biology, reveals that morphogenesis of the basally invaginating OC is initiated by active patterned cell behaviours at the apical surface.

Engaging in a constant feedback loop between experimental findings and theoretical modelling allowed us to generate new insights and ideas for how the eye acquires its hemispheric 3D shape. We reveal that in-plane cellular rearrangements at the apical surface of the NE generate the spontaneous strains that initiate invagination of the NE basal surface. Concurrently with these cell rearrangements, another set of patterned cell behaviours at the apical surface of the RPE leads to the spontaneous strains that drive the curvature of this epithelial layer. While these two active patterns evolve independently from each other, the cooperation between them is important for sculpting the final shape of the OC.

Our work thus reveals an unexpected mechanism to drive OC shape emergence, highlighting a central role for the apical surfaces, which have previously not been explored. We show that in-plane patterned cell behaviours are sufficient to drive 3D shape transitions in a complex structure like the OC. We also show that independent patterns of cell behaviour can work together to achieve a single developmental outcome. These concepts can potentially be applied to the study of OC morphogenesis in other vertebrates, but also to other complex, multilayered structures, such as the growth of the zebrafish pectoral fin^45^, shape dynamics in cnidarians^46^ and many more.

During OC formation, discrete morphogenetic events, such as rim cell migration, RPE cell flattening and NE cell shape changes occur in parallel in different regions of the forming cup. Together, these coupled, concurrent behaviours make OC shape generation a highly complex process. To start deconstructing this complexity, we focused on the initial key step of overall OC formation: the onset of OV invagination. While NE basal acto-myosin activity, rim cell migration and lens formation contribute to overall OC morphogenesis later in the process as established previously^14,18–22^, our findings show that they do not drive the *onset* of invagination.

We show that OV invagination starts at the apical surfaces of the RPE and NE. While strains originated at the apical surface have been previously proposed to be important for invagination progression^27^, theoretical approaches for *ex-vivo* generated retinal organoids focused on modulation of the effective material properties by local thinning of the tissue in systems with significant luminal volume. However, these organoids develop in different conditions so that it is likely that other mechanisms are at play *in vivo*.

In this *in vivo* study, we take a different approach using inspiration from shape programming frameworks and combining them with theoretical modelling and experiments, we reveal active spontaneous strain pattern candidates that can lead to RPE and NE apical surface curvature transitions. In the RPE, a pattern of cell flattening creates a gradient of apical cell area growth that leads to the cupping of the RPE layer. While at the NE apical surface, we show that cells exchange their apical neighbours consistent with an elliptical pattern. We propose that these patterned rearrangements cause the emergence of the spontaneous strains that lead to a concurrent cupping of the NE apical surface.

While the two apical surfaces curve independently of each other, they start this process at approximately the same time. This coordination potentially contributes to the overall OC shape in at least two ways: 1) by ensuring that the RPE correctly wraps around the future retina and 2) by ensuring that the NE has the necessary space to invaginate. It remains to be seen how exactly this coordination is achieved. Given that we found that the RPE and the NE remain connected through their apical sides, one possibility is that this connection provides a mechanical feedback mechanism that is able to synchronize the activity on these two surfaces. Such mechanical feedback could also contribute to the formation of the correct optic cup shape. However, while the connection between the two apical surfaces keeps the two epithelia together, it also needs to remain flexible and dynamic enough to allow for distinct cell movements. While these insights are compelling, further work is needed to understand these apical-apical connections, including their molecular identity, how they are regulated, their overall dynamics, and their possible roles throughout late eye development. In the future, shedding light on what keeps these two epithelia together will contribute to the knowledge of OC formation, while potentially also bringing new insights about the morphogenesis of apposing epithelia, such as the drosophila wing disk or the vertebrate neural tube.

Our theoretical model shows that the emergence of curvature at the apical surface is sufficient to drive invagination of a more “passive” basal surface. Indeed, when we experimentally disrupted patterned rearrangements at the apical surface, the NE basal surface failed to initiate its invagination. Given that shape transitions lead by activity at the apical surface are typically associated with apical constriction, our work supports a new morphogenetic mechanism for the emergence of 3D shapes driven by the apical surface. At this point, it is not fully clear how the strains that initially occur at the apical surface are transmitted to the NE basal surface, leading to its invagination. One possibility is that cell rearrangements at the NE apical surface are transduced down the lateral sides of the cells to the basal surface on a slower timescale, dependent on the height of the epithelium. While cell rearrangements are common during morphogenesis (for example the convergence-extension during tissue elongation^47^), their role in 3D shape transitions has not been fully explored^10^. Apical-to-basal topological transitions (AB-T1s) have been shown to occur on curved, thick epithelial monolayers, as a way for the cells to adapt to the geometry of the tissue^48^. Taking our results into account, it is possible that these previously described cell rearrangements at the apical surface are not only passively induced by tissue curvature, but that they, in fact, can drive the curvature of the tissue themselves.

While our study focuses on the physical mechanisms that drive curvature onset, the specific roles of the other concurrent morphogenetic events to the final OC shape remain to be integrated into our findings. For example, NE basal constriction^18^ could be necessary for the cells to adapt to the changing geometry of the tissue. Another interesting question is how the curvature of the lens is coordinated to fit into the cavity formed upon invagination. The fact the NE basal surface curvature increases upon lens cell ablation points to the existence of a pulling force from the NE to the lens. Given that several studies have reported the existence of filopodia connecting these two tissues, it is possible that a mechanism exists by which the NE pulls the lens into shape. Interestingly, when the basal surface fails to invaginate upon Shroom4 disruption, presumptive lens cells accumulate between the OC and the ectoderm, without forming their usual round shape (Fig. 5c). While we cannot exclude that the PCP pathway is also involved in lens formation, it is possible that, in the absence of an OC cavity, the lens fails to round up due to the lack of space or the absence of a pulling force from the NE. Narrowing down which processes have an active vs passive role throughout OC formation, will ultimately allow for an even better understanding of OC morphogenesis.

Overall, our comprehensive 3D shape analysis and modelling approach holds the potential to expand our understanding of how patterned cell behaviours lead to shape transitions in diverse contexts. Beyond formation of the OC, we shed light on a new mechanism leading to 3D shape acquisition, that relies on active, in-plane, collective cell behaviours and on the cooperative reinforcement of two active patterns to drive a target shape. Ultimately, our approach is a significant step towards a more complete understanding of 3D shape acquisition, a long-standing question in Biology and Physics.

## Methods

### Zebrafish husbandry

Wild-type zebrafish and transgenic lines (Danio rerio; AB and TL strains) were maintained at 28 °C. Embryos of yet undetermined sex, between 6 and 22 somite stage^49^, were raised at 21ºC or at 28.5ºC in E3 medium. All animal work was conducted in accordance with institutional standard operating procedures under the licensing of the DGAV (Direcção Geral de Alimentação e Veterinária, Portugal and in accordance with the European Union directive 2010/63/EU and with the Portuguese Decree Law nº113/2013.

### Transgenic lines

The following stable transgenic lines were used: tg(lama2:lama-sfGFP); tg(lama2:lama-mKate2)^21^ – to visualize the extracellular matrix; Tg(ß-actin: mNeonGreen-zo-1b)^50^ – to label the apical surface; tg(RX2:GFPcaax)^22^ – to visualise the optic vesicle and optic cup epithelium; Tg(h2az2a:h2az2a-GFP) ^51^- to label all the nuclei; tg(ß-actin: Ras-GFP) ^52^ – to label cell membranes; tg(ß-actin:Actin-CB-GFP), this study– anti-actin nanobody to quantify actin intensities.

### mRNA injections

Embryos were injected with 1nl of a 25-50ng/µl mRNA solution at the 1-2 cell stage to label all cells of the embryo or between 64-128 cell stage for sparse/mosaic labelling. RNA transcription was performed with an Ambion mMessage mMachine kit. The following templates were used: pCS2+ GFP-zZO-1b ^50^ (kind gift from CP lab) and pCS2+ mKAte-zZO1b (this study) – to label apical surface ; pCS2+ Utrophin-GFP^53^ – to label utrophin; pCS2+ MRLC2-GFP^54^ – to label myosin; pCS2+ Dsh-DEP+ ^36,37^ – Dominant negative form of Disheveled (kind gift from the Barriga lab).

### Plasmid construction and generation of transgenic lines

The T2-ß-actin:ActinCB-GFP plasmid was assembled using Gateway cloning (Thermo Fisher Scientific) based on the Tol2 kit^55^. Briefly, the Actin-VHH-GFP (Actin Chromobody, ChromoTek)^56,57^ was isolated from the plasmid tol2-UAS:Actin-CB,crybb:eCFP (kind gift from P. Panza) by PCR, to create a middle entry clone (pME ActinCB L1L2), using the following primers: 5’ggggACAAGTTTGTACAAAAAAGCAGGCTtcacc3’ and 5’gggggACCACTTTGTACAAGAAAGCTGGGtcttac3’ ; The pME was then recombined with the plasmids b-actin promotor_p5E (L4-R1) and pTol2+pA_pDEST(R4-R2) with LR clonase plus to create the final plasmid.

Using this plasmid we created the stable transgenic line tg(ß-actin:Actin-CB-GFP). To this end, one cell stage wildtype embryos were injected with 1nl of a solution containing the plasmid T2-ß-actin:ActinCB-GFP at 20ng/µl, plus Tol2 transposase RNA at 50ng/µl, in ddH_2_O. F0 embryos were selected for GFP expression at 24hpf and raised. Germline carriers were identified by crossing the F0 generation with wildtype fish.

The pCS2+ mKAte-zZO1b plasmid was assembled using the Gibson Assembly cloning Kit (NEB), in a reaction containing three fragments: the pCS2+ backbone, the mKate fragment and the the zZO1b fragment. The pCS2+ backbone fragment was obtained by digesting the pCS2+ plasmid with XbaI and BamHI; the mKate fragment was obtain by PCR from the Evrogen plasmid pmKATE2, using the following primers: 5’-CTTGTTCTTTTTGCAGGATCGATCCACCGGTCGCCACCAT-3’ and 5’-attcgagatctgagtccggaTCTGTGCCCCAGTTTGCTAG-3’; the zZO1b fragment was obtained by PCR from the plasmid pCS2+ GFP-zZO-1b ^50^, using the following primers: 5’-CTAGCAAACTGGGGCACAGAtccggactcagatctcgaat -3’ and 5’-ACGACTCACTATAGTTCTAGTCAGAAGTGGTCGATCAGCA -3’.

### Morpholino injection

2ng of Shroom4 morpholino (ACATTTGTGTGTTTGCTTACCTTCG)^44^ were injected together with ZO1-mKate2 mRNA, into tg(lama2:lama-mKate2) one cell stage embryos.

### Imaging

For live-imaging, embryos were mounted in 0.8% low melting point agarose, in mattek glass bottom dishes and kept at 28,5ºC throughout imaging. Image acquisition was carried out either on a Zeiss LSM 980 (using multiplex Airyscan 8y), using a 25x 0.8NA multi-immersion objective, or on a Andor DragonFly SDC using a 40x 1.15NA water/silicone objective, or on a 3i Marianas SDC using a 40x 1.15NA water/silicone objective.

For the 3D reconstruction of OC morphogenesis, embryos were imaged on a Zeiss Z1 lightsheet system, using a 20x 1.2NA water immersion objective

### Laser ablation

Laser ablation was carried out on a 3i Marianas SDC equipped with a 3i Ablate! Laser ablation system, using a 355nm pulsed laser. Prior to ablation, laser position was calibrated using fluorescent agarose, to mimic the mounting conditions of the embryo. Ablation was performed using a 20x or 40x water objective. Imaging was carried out before, during and after ablation.

### Segmentation

The ECM signal of tg(lama2:lama-FP) embryos outlining the optic vesicle to optic cup was segmented using the LimeSeg plugin for Fiji^58,59^ and the skeletonseg option, together with a customised script for automating the segmentation over time (supl Fig 2a). Segmentation results were saved as a separated PLY mesh for each time point. Meshes were visually assessed and, if needed, further processed on Blender (www.blender.org) to correct segmentation errors and/or to remove optic stalk remnants.

To segment the apical surface, we first obtained a probability map of the ZO1-FP signal by running a pixel classification with Ilastik^60^. Next, the ‘find edges’ filter on Fiji was used to process the probability map. The resulting image was then segmented with LimeSeg using the sphereSeg option. Segmentation results were saved as a separated PLY mesh for each time point.

Final PLY meshes were processed with a customised Python script to extract geometric properties and for the analysis of curvature progression (details bellow).

Nuclei Segmentation was performed with StarDist, using a previously published trained model^28,61^. To count the nuclei that belonged to the Optic Cup only, ECM segmentation was combined with the nuclei segmentation results by using a customized Python script. First, the ECM segmentation was converted into a mask file; this mask was then overlapped with the segmented nuclei and all the nuclei outside were excluded (Extended data Fig. 2b).

### Mesh analysis and region separation

We developed a computational pipeline to analyse the spatiotemporal changes in the geometry of the optic cup using Python-based tools (Extended data Fig. 2d-d’). Meshes obtained from the LimeSeg segmentation were processed using PyMeshLab^62^ to generate manifold surfaces with closed boundaries.

For each mesh vertex, we calculated several geometric properties: mean and Gaussian curvatures [PyMeshLab curvature estimation], distance from the structure border, normal angle relative to the horizontal plane, and distance from the fitted midplane. These features then served as the basis for segmenting the structures into distinct regions. Initially, Gaussian Mixture Models [scikit-learn] classified the mesh vertices into three clusters based on their geometric properties, separating border regions (corresponding to the rim region) from non-border regions. Subsequently, DBSCAN clustering [scikit-learn] was used to refine this classification by ensuring spatial continuity of the retinal pigmented epithelium and neuroepithelium surfaces.

For analysis of surface properties, we quantified the distribution of mean and Gaussian curvatures in the segmented regions, over sequential timepoints.

### Fluorescence intensity measurements

A schematic representation of the image analysis pipeline to quantify fluorescence intensity can be found in Extended Data Fig. 2c. For actin and myosin fluorescence intensity quantification a 2D segmentation, at the centre of the OC, of the ECM signal was performed using the Kappa plugin for Fiji^63^ to fit a B-spline curve onto the lama-FP signal. This was used to represent the outline of the OC over time. This curve was further used to retrieve point curvature data. Spatial curvature representations over time and respective kymographs were then obtained in Matlab 2018a/2020a, with further plotting and analysis in Python for actin and curvature.

Based on the 2D segmentation, the OC was divided to represent four distinct regions: two rim areas at nasal and temporal pole, the neuroepithelium and the retinal pigmented epithelium. A second division was created by dividing the full optic cup space into five layers, from the optic cup outline to the inside. Here the first four layers have the same imposed thickness (3.7 µm). Pixel intensity was then retrieved within each combined portion and normalized to the overall optic cup intensity.

### Simulations

#### Spring lattice mechanics

We use a programmable spring lattice model in two distinct geometries. The first is a simplified elliptically symmetric 4 layered mesh, and the second is an effective mesh generated from the experimentally derived segmentations of the distinct OC layers. In both cases, the lattice edges are overdamped elastic springs with programmable spring constants. The elastic energy of the entire lattice is given by

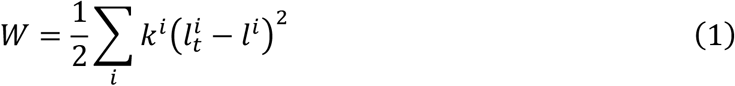

Where *i* denotes a single spring; *k*^*i*^ is that spring’s constant; 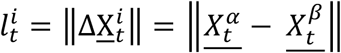 is the length of spring *i* at time *t* given by the positions of the vertices *α, β* that are connected by that spring; *l*^*i*^ is the corresponding’s spring rest length. At *t* = 0 the model is stress free, meaning all the springs’ rest lengths are equal to their initial lengths, so

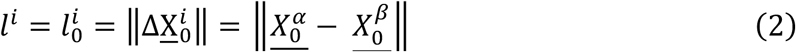

The model tries to find a preferred configuration of vertex positions by minimizing *W* over time after a new pattern of rest lengths are prescribed.

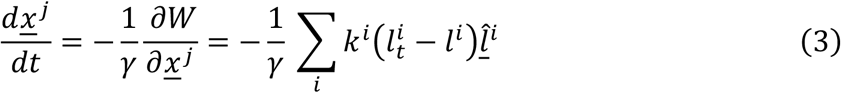

Here *<u>x</u>*^*j*^ is the current position of vertex *j*. The sum is over all the springs *i* connected to vertex *j* and 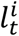 the length of the spring. 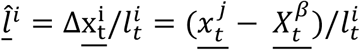 is the unit vector along the spring *i* connecting vertex *j* and *β*.

#### Shape programming through spontaneous strain tensor

In this work, we model the OC invagination as a shape change that arises from spontaneous strains, a change in the ground state of the local length scales of the different layers. We realize this using a spontaneous deformation gradient tensor field, 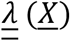, a rank 2 tensor field. Each component in 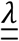 is a multiplicative factor by which the rest lengths of the tissue change in a given direction. Generally, in an arbitrary coordinate system, 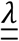 can be written as

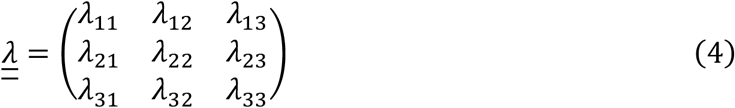

We choose the coordinate system so that it aligns with our deformation pattern, specifically we chose basis vectors *<u>e</u>*^1^ and *<u>e</u>*^2^ tangent to the surface and aligned with the principal axes of in-plane deformation, and *<u>e</u>*^3^ the surface normal. In this case, 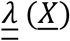 has a diagonal form

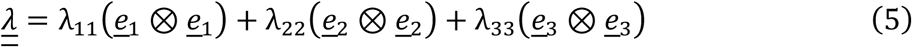

Furthermore, the surface components *λ*_11_, *λ*_22_, can be decomposed into their isotropic and anisotropic components. Isotropic deformations change the local area of the surface by changing the local rest lengths equally in all directions. Anisotropic deformations increase the local length in one direction and decrease the local length in the other direction while preserving the local area. Hence, we may write the surface components as follows:

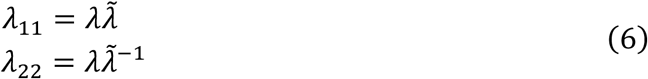

where *λ* is the magnitude of isotropic deformation and 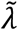 is the magnitude of anisotropic deformation. In this work, each layer may experience a different spontaneous strain field corresponding to the biological activity in that layer. Hence for each layer a coordinate system was chosen as described and the layer’s spontaneous deformation tensor can be written as

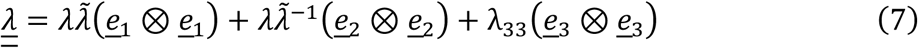

#### Computational realization of the model in TopoSPAM

To implement our model computationally, we use TopoSPAM, a python simulation platform for morphogenesis and biological active matter^64^. TopoSPAM enables creating a spring lattice using a given mesh, applying different spontaneous strain patterns to it and simulating the resulting shape change caused by that pattern. To achieve that the spontaneous strain field, 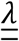, is discretized in the following manner. For each spring 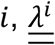 is the average value of the spontaneous deformation of the vertices *α, β* that it connects

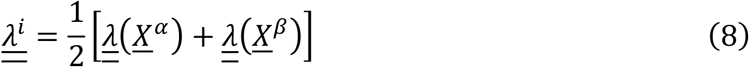

Next, each spring’s rest length is updated to reflect the effects of the spontaneous deformation field, for an initial spring rest length 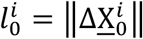 we update it as follows

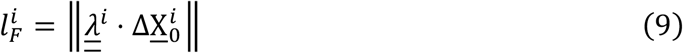

Finally, TopoSPAM relaxes the model using Equation 3 by updating the position of the vertices with

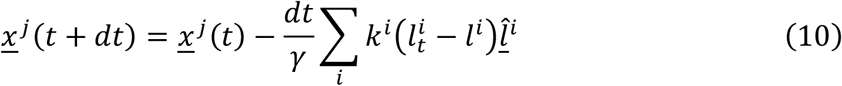

The relaxation continues until the mean movement of the vertices in a time stop falls below a given threshold, here, 2 × 10^−6^.

#### Generating the simplified double layered mesh

To test the three different potential mechanisms of shape change, we generated a simplified mesh to represent the apical surfaces of the OC. We started with TopoSPAM’s circular mesh, a triangulated network of two flat circles connected to one another. As the OC has a complex shape that isn’t radially symmetric, but instead more closely resembling an ellipse, we need to deform the mesh to better capture the boundary conditions of the OC. We first deform the circle to an ellipse with an eccentricity that mimics the experimentally extracted apical surfaces’ eccentricity. Next, we choose, *d*_*xapical*_, the separation between the two layers such that the ratio between the major axis length, *a* and the separation *d*_*xapical*_ in the simplified mesh matches the experimentally extracted parameters. Here we choose the in-layer springs constant *k*_*apical*_ = 1 and the springs connecting the layers to have *k*_*xapical*_ = 0.7. Here, we exclude the rim region of the OC from all following meshes. While including the rim might enhance the coupling between the layers and add to the mechanical stability of the system, it also adds several parameters to the model regarding the patterns behaviour and matching in those regions. As we don’t have experimental data to inform us about these, we chose to exclude the rim region and focus on the bulk of the OC.

#### Simple mechanisms of cupping

Using the simplified double layered mesh, we apply three different basic spontaneous strain patterns that can drive cupping of flat surfaces. In these simulations the spontaneous strain field was applied only to the apical in-layer springs, while the springs connecting the apical layers remain at rest. The first is global layer area mismatch, here we apply a uniform in-plane isotropic growth to an entire layer. To achieve an area mismatch, we apply different growth magnitude to each layer. In the form of Equation (7) we apply a spontaneous displacement field of the form to layer *l*

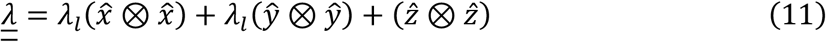

Here *λ*_*l*_ is the isotropic growth magnitude applied in-plane of each layer with *λ*_*RPE*_ > *λ*_*NE*_. In our simulation we chose *λ*_*RPE*_ = 1.25; *λ*_*NE*_ = 1.2.

The second is radial gradient isotropic growth^65^. This takes the form

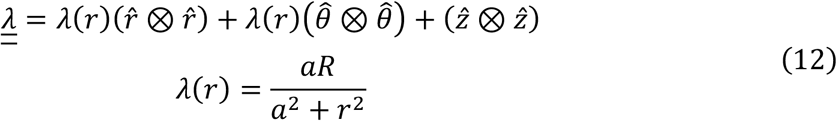

Here *R* is the target radius of the cap and 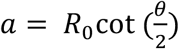 with *θ* the cap section angle and *R*_0_ the initial radius of the flat surface. We apply the same pattern to both layers with *a* = 1.2; *R* = 1.5.

The third is a uniform radial anisotropic pattern which takes the form

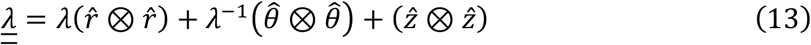

Here we choose *λ* = 1.1 so at each point the material tries to extend along the radial axis and contract tangentially to it. As in the second case, we apply the same pattern to both layers. These all are able to drive the simplified double layered mesh to form a cup. As both the gradient of isotropic growth and the anisotropic pattern are symmetric about the z-axis, there are solutions of cupping in both the positive and negative z-direction. This may lead to final configurations that are a mix of both the cups. To avoid these and break the symmetry in our simulations we introduce a small initial curvature *κ* = 0.05 to the stress-free configuration. Unless otherwise stated, all other simulations with the simplified mesh use a completely flat initial configuration.

#### Ablation simulation

To achieve the effect of the ablation experiment, we increase the separation distance *d* between the apical surfaces to 4*d*_0_ which correlates to the observed increase in that distance in the experiment as well as decreasing the apical connecting springs *k*_*xapical*_ = 1 × 10^ − 5 to simulate the severing of the connection between the layers. As TopoSPAM does not explicitly define a bending rigidity in the model, this arises as a result of the thickness of the mesh, which is *d* in our case. As we effectively disabled the connection between the layers, each layer acts as an independent 2d surface with a pattern on it, leading to numerical instabilities that result in final configurations that can be very noisy and exhibit multiple self-intersections. To overcome this, we double each of the apical layers with an exact copy of itself, experiencing the exact same spontaneous deformation pattern and separated by distance 0.5*d*_0_ from its original layer. This serves to add a bending rigidity to each layer, stabilizing the simulation and allowing us to capture the effect of the ablation on the simplified geometry. We than apply the same strain patterns as described above to the new ablated mesh. Here again we introduce the same small initial curvature for the gradient of isotropic growth and the anisotropic pattern simulations to break the initial symmetry as described above.

#### Generating the basal height map

During flattening, RPE cells simultaneously significantly increase their in-surface area while decreasing their height. To gain insight into the potential patterned flattening of the RPE, we use the experimentally segmented apical and basal surfaces to calculate the vertical separation between the two. This serves as a proxy for cell height, as the cell height and cell area are anti-correlated due to the conservation of cell volume. Hence, a patterned decrease in height is accompanied by a patterned increase in cell areas. We calculate the vertical separation between the surface by looking at the effective networks we generate from them and then finding for each vertex on the basal side the closest vertex in the apical side. We generate a height map for each experimental time point to create the time-series dataset of height maps.

#### Semi-Confocal Elliptic Coordinate System

We observed that RPE flattening in the OC initiates further from the neural stalk, aligning with the approximate focus of its enclosing ellipse. To accurately model this spatial characteristic and integrate it with the boundary conditions, we adopted a specialized coordinate system. This semi-confocal elliptic coordinate system defines a family of encapsulating ellipses sharing one fixed focus, defined by the following equation

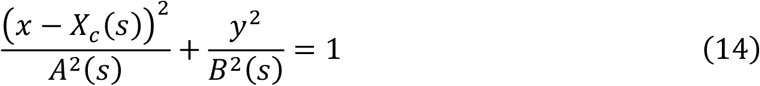

With 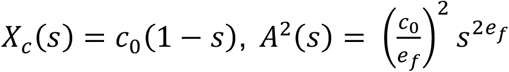 and 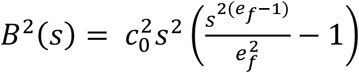. Here, *s* ∈ [0,1] is the ellipse shape parameter; *c*_0_ defines the fixed position of the shared focus (*c*_0_, 0) in the coordinate system of the mesh; and *e*_*f*_ is the target eccentricity of the ellipse when *s* = 1, used to match the boundary conditions. Next, we define a 3D curvilinear coordinate system with local orthogonal basis vectors derived from Equation 14, as follows:

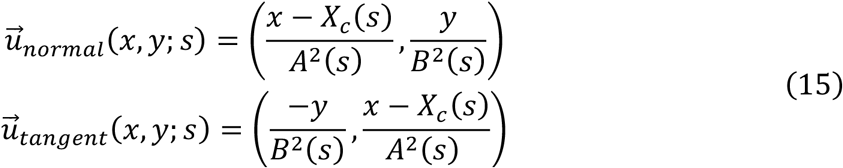

So the principal directions are given by

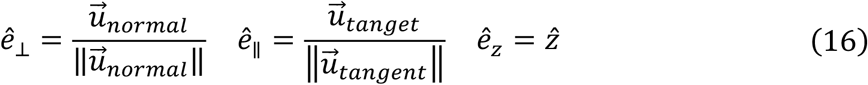

In the following sections, we obtain s from the Cartesian coordinates of the mesh vertices. We then use this s value to apply the different patterns to define the correct pattern shape within this semi-confocal elliptic coordinate system. A full derivation of this equation is provided in the Supplementary Material.

#### Experimentally informed spontaneous strain patterns

To implement our hypothesised experimentally informed spontaneous strain pattern we apply a different pattern to each of the apical surfaces. To capture the gradient of RPE flattening, we applied a gradient of isotropic growth in the semi-confocal elliptic coordinates to the RPE layer in a similar fashion to Equation (12):

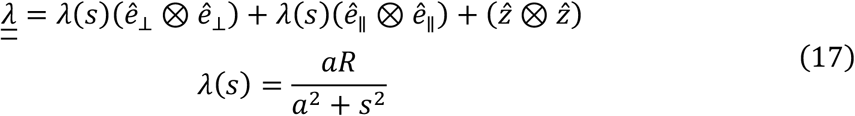

To capture the observed neighbour exchanges on the NE that exhibit a bias to contracting along the OC major axis, we apply an anisotropic spontaneous strain^66^ in an elliptic coordinate system to the NE layer where *ê*_*μ*_ and *ê*_*v*_ are the principal directions along the ellipses and the hyperbolae respectively. To match the NE pattern center of invagination with the RPE one, we add a bias in the magnitude using the semi-confocal ellipse parameter, *s*, as follows:

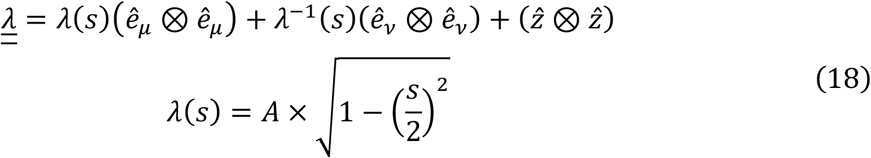

#### Simulating the isolated apical surfaces

To simulate the effect of the layer-specific strain pattern above on the apical surfaces we use our simplified elliptical mesh. We choose the in-plane apical spring constants, *k*_*apical*_ = 1, to be stiffer than the springs connecting the apical layers, *k*_*xapical*_ = 0.7. It is important to note that the springs connecting the two apical layers are an effective description of both the thickness of the apical layers and the biological connectors between the apical surfaces. This is because in the experimental data we are not able to resolve each apical layer separately and the segmentation of the apical layers is assumed to be a mid-plane. Due to this fact the apical cross-connecting springs rest lengths are updated to capture a combination of the activity inside the layers as well as the assumed inactivity of the apical cross-connectors. This is achieved by averaging the calculated new rest lengths resulting from the RPE layer’s isotropic pattern, the NE layer’s anisotropic pattern, and the original at-rest rest length of these springs. For the distinct patterns we choose the following parameters *a* = 4, *R* = 4.4, *A* = 1.3.

#### Generating the full simplified mesh

To generate a simplified mesh of the OC we extend the apical simplified mesh by adding the basal surfaces as copies of each apical surface. We fully connect the layers and choose the separation distance between each layer’s apical and basal surface, *d*_*layer*_, such that the ratio 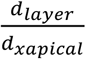 in the simplified mesh matches the ratio of the segmented layers from the experimental data. We again set *k*_*apical*_ = 1 and *k*_*xapical*_ = 0.7. Next, we set the springs of the basal surfaces, *k*_*basal*_ = 0.05, to be the softest, as we assume these are the softest in the biological system. Finally, we set the springs that represent the lateral stiffness of the cells *k*_*NE*_ = 0.5, *k*_*RPE*_ = 0.5. The lateral sides are chosen to be relatively stiff as during OC invagination the NE cells do not change their heights significantly. We assume that the significant change in the heights of the RPE cells is actively driven.

#### Simulating the full simplified OC

To simulate the effect of our hypothesised layer-specific strain pattern on the full simplified mesh we apply to the mesh similarly to the isolated apical surface case with several additions. As the RPE area change is assumed to not be localized to the apical side, we apply the same pattern given by Equation 17 to the RPE basal side. Furthermore, as the RPE lateral is undergoing significant decrease in size, we apply a uniform contraction along the 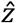 direction of the springs connecting the RPE basal and apical surfaces

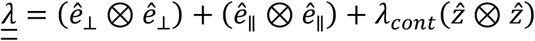

Here, *λ*_*cont*_ =< *λ*(*s*) >^−2^ is the inverse square mean of the isotropic growth of the basal surfaces given by Equation 17. The NE lateral and basal springs are kept at rest as we assume the initial anisotropic pattern is localized to the apical surface. The results in the manuscript arise from the following parameter choice *a* = 4, *R* = 5, *A* = 1.3 and are representative of the parameter regime that leads to correct initial invagination of the mesh. To further analyse the parameter space, we ran a sweep of the model’s pattern parameters: *a* ∈ [1.5, 1.89, 2.28, 2.67, 3.06, 3.44, 3.83, 4.22, 4.61, 5], *R* ∈ [2.5, 2.78, 3.06, 3.33, 3.61, 3.89, 4.17, 4.44, 4.72, 5], *A* ∈ [1, 1.04, 1.09, 1.13,1.18,1.22, 1.27, 1.31, 1.36, 1.4]. The results show a wide range of parameters in which a strong cup forms as well as zones of weak cup formation, numerically unstable results and a zone of RPE lateral extension caused by an average RPE local area shrinkage (Extended Data Fig. 7). Finally, to simulate the experimental perturbation of the apical surfaces, we run the simulation with no pattern on the NE apical surface (Extended Data Fig. 8), so all the NE apical layer’s springs are left at their initial stress-free state. We maintain the pattern on the RPE layer as described above.

## Acknowledgements

The authors would like to thank the Cell Biology of Tissue Morphogenesis laboratory for ongoing input and project discussion.

João Coelho and Renata Cunha are thanked for help with cloning, Lucrezia Ferme for establishing the nuclear segmentation and filtering pipelines, Bruno Vellutini and Byung Ho Lee for sharing image analysis scripts. We further thank the Bioimaging Facility and Aquatic facilities at the Gulbenkian Institute for Molecular Medicine (GIMM, formerly Instituto Gulbenkian de Ciência, IGC) for experimental and technical support. The authors would further like to acknowledge Elias Barriga for discussions, kind gift of reagents and support at later stages of writing. Juan Martinez-Morales, Pierre Haas and Stephan Grill are thanked their helpful comments on the manuscript,

Caren Norden discloses support for the research of this work from European Research Council (ERC) consolidator grant (H2020 ERC-2018-CoG-81904). Ana Patricia Ramos was supported by Marie Skłodowska-Curie Actions (101038054). Alicia Szalapak was supported by the European Union’s H2020 research and innovation programme 6 (829010 (PRIME H2020-FETOPEN-2018-2019-2020-01)).

## Author contributions

CN: conceptualization, funding acquisition, supervision, writing – original draft;

CDM: conceptualization, methodology (theoretical model), funding acquisition, supervision, writing – original draft;

APR: conceptualization, methodology, experiments, data acquisition, quantification and analysis, funding acquisition, writing – original draft and figures;

LM: Theoretical simulations, writing – original draft;

AS: mesh processing and analysis pipelines;

LD: protein quantification pipelines (with input from APR), formal analysis;

MH: formal analysis

## Extended data figures and legends

**Extended Fig. 1.**
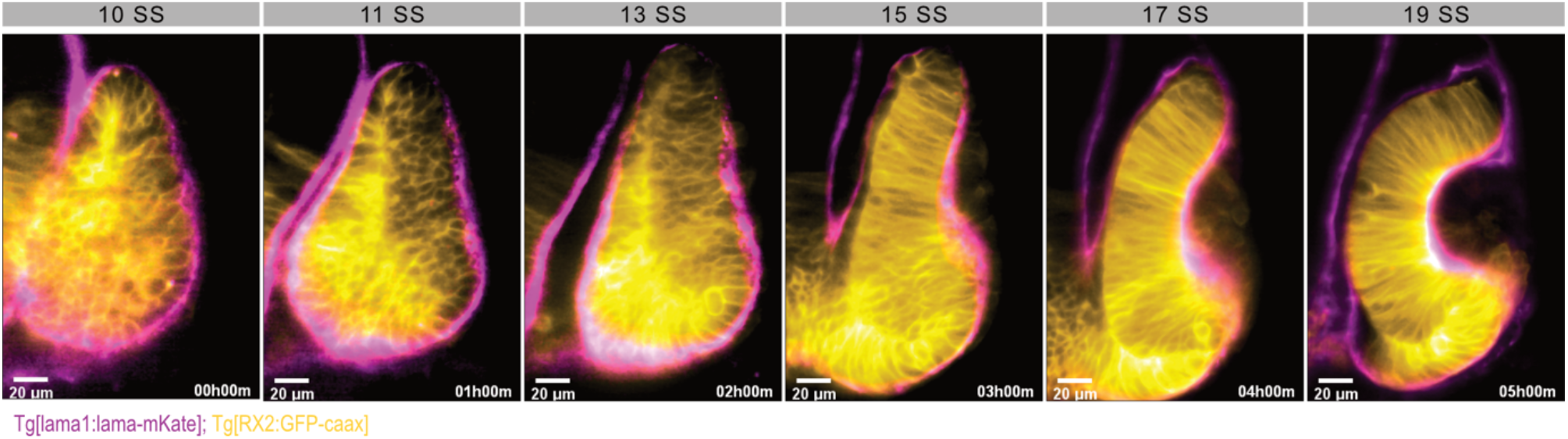
Extracellular Matrix Follows Optic Cup shape changes. Lightsheet optical slice of the developing OC. Basal surface (ECM) is labelled with lama-mKate (magenta), and the membranes of OC cells are labelled with GFP-Caax, driven by the OC specific promoter, RX2 (yellow). Developmental time is indicated in somite stage (SS)

**Extended Fig. 2.**
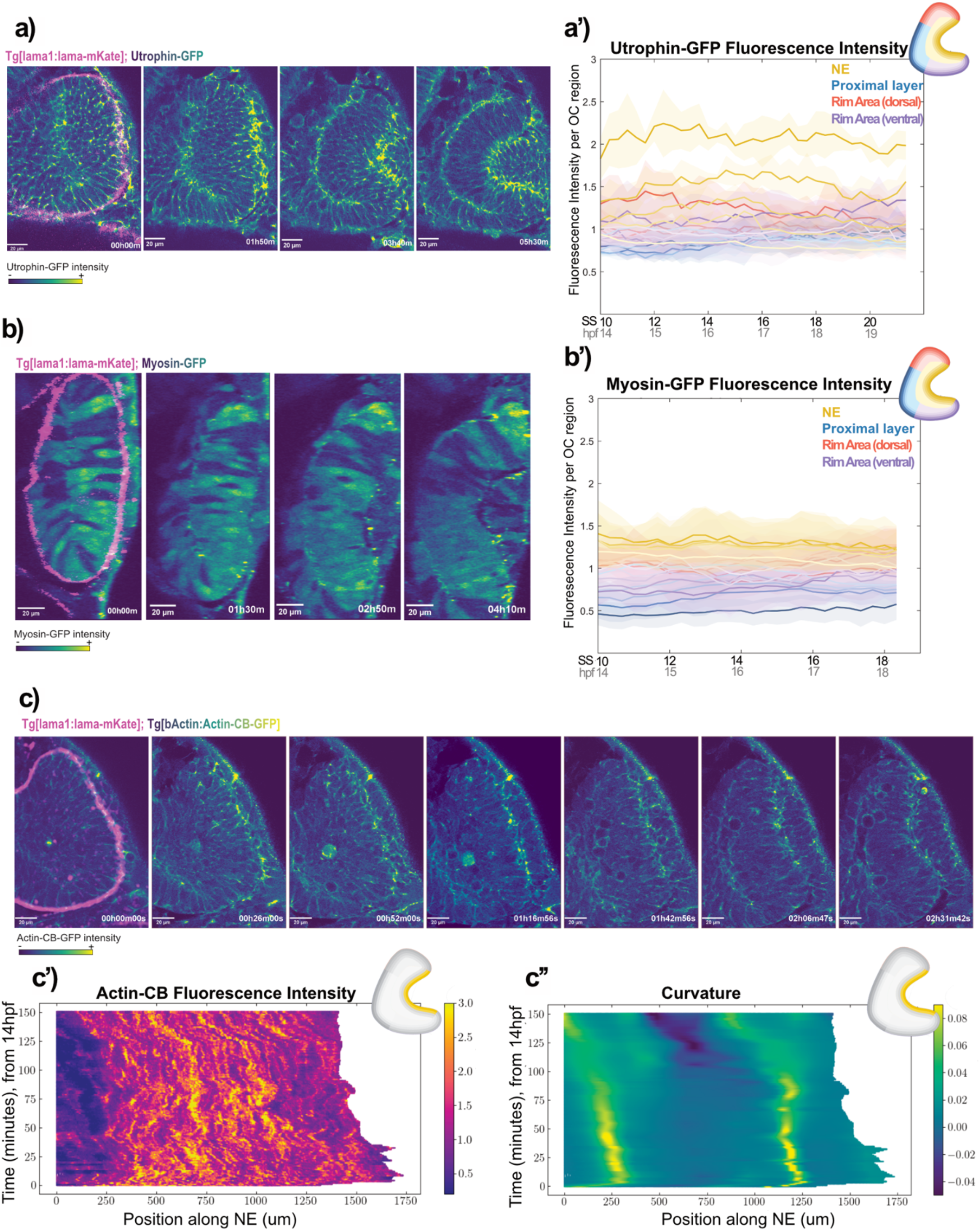
Acto-Myosin protein quantification during Optic Cup development. **a**, utrophin quantification, AiryScan optical sections. The basal surface (ECM) is labelled with lama-mKate (magenta), Utrophin-GFP intensity is depicted using a sequential colour map; **a’** Fluorescence intensity quantification of the Utrophin-GFP signal at the basal surface of all the regions of the OC, over time. The inlet with optic cup scheme represents the colour code for the different regions.**b**, myosin quantification, AiryScan optical sections. The basal surface (ECM) is labelled with lama-mKate (magenta), Myosin-GFP intensity is depicted with a sequential colour map; **b’** Fluorescence intensity quantification of the Myosin-GFP signal at the basal surface of all the regions of the OC, over time. Inlet with optic cup scheme represents the colour code for the different regions.**c**, actin quantification, AiryScan optical sections. The basal surface (ECM) is labelled with lama-mKate (magenta), ActinCromoBody-GFP intensity is depicted with a sequential color map; **c’** Histogram with fluorescence intensity quantification, over time, of the ActinCB-GFP signal at the basal surface of the NE only, as represented in yellow in the optic cup scheme; **c’’** Histogram with mean curvature quantification along the same NE region as in **(c’)**

**Extended Fig. 3.**
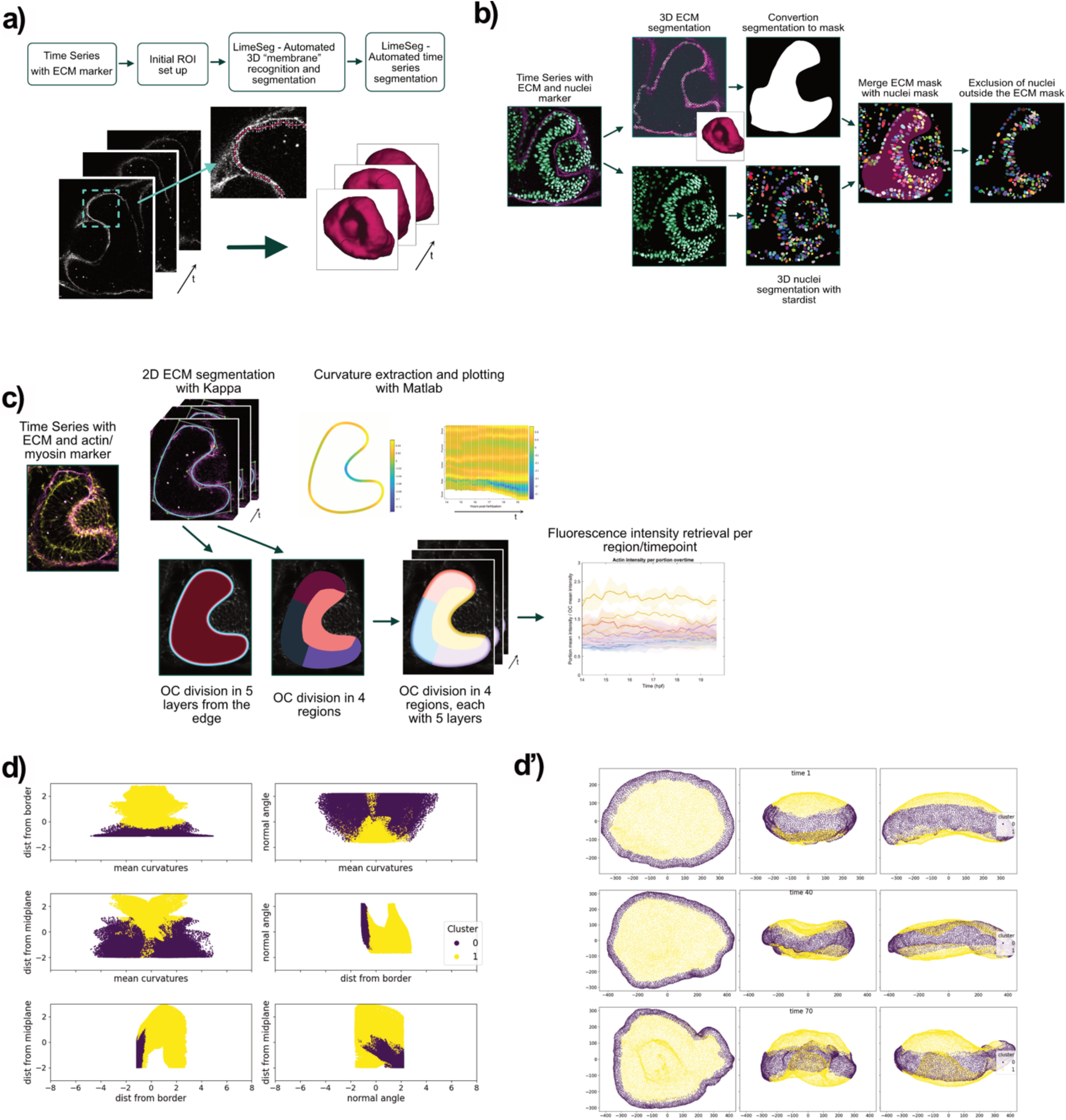
Image analysis pipelines. **a**, segmentation of the extracellular matrix during optic cup development, using LimeSeg**; b**, optic cup nuclei segmentation using stardist. **c**, segmentation and partitioning of the optic cup and extraction of fluorescence intensity values in the different regions, during optic cup formation; **d-d’**, mesh processing pipeline to separate the optic cup in three regions, **b**, vertex properties analysis and separation into 2 clusters, **d’** distribution of the vertex clusters in 3D, in 3 representative timepoints.

**Extended Fig. 4.**
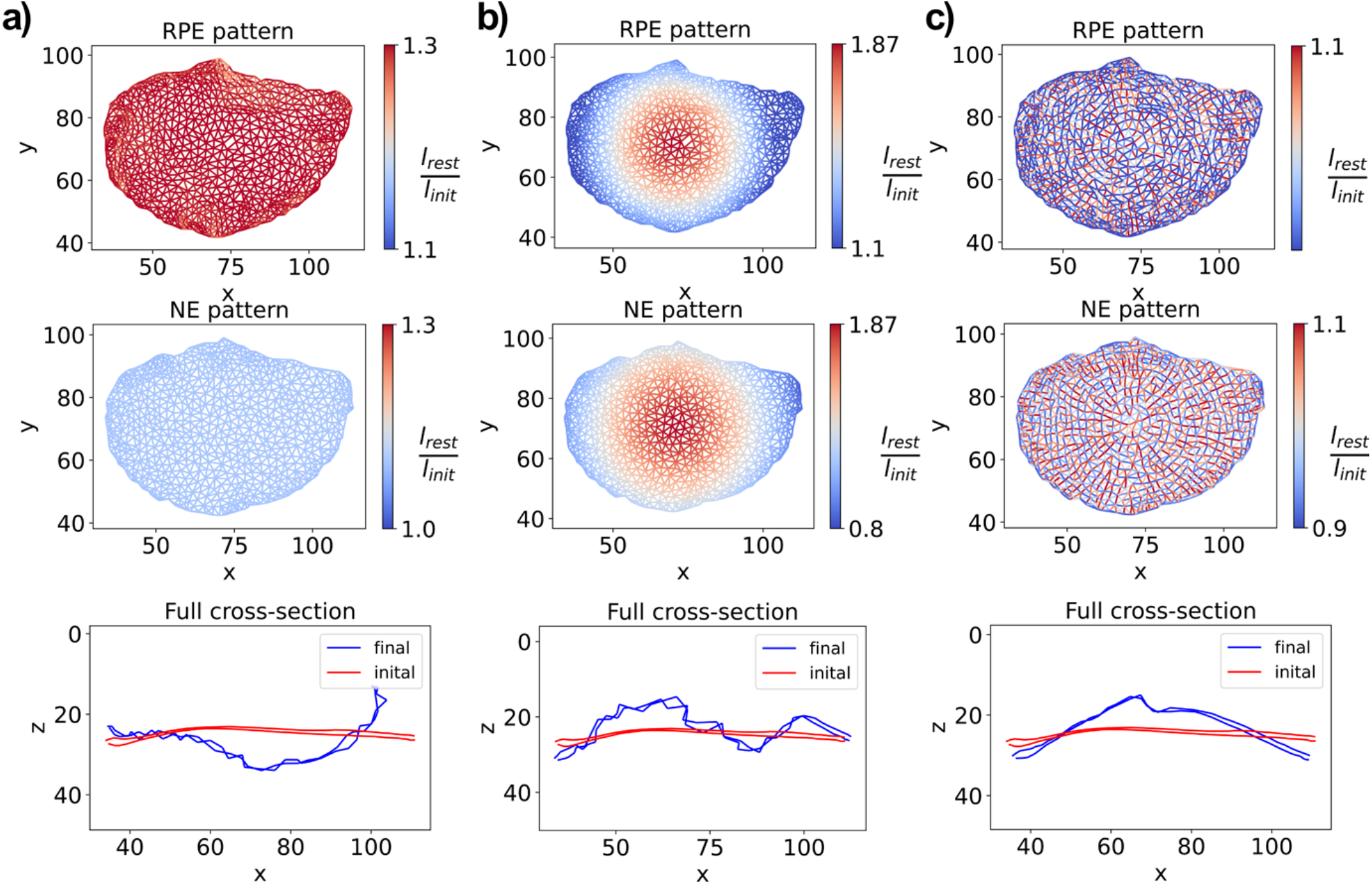
Simulations of the theoretical candidate patterns for RPE and NE apical surface cupping on segmented apical meshes. The three theoretical candidate patterns applied to the segmented apical surfaces from the experimental data segmentation (z-axis flipped to maintain orientation convention as RPE layer is above the NE layer in the extracted mesh). **a-a’**, strain pattern resulting from a total increase in cell area at both layers, here *λ*_*NE*_ = 1.1 and *λ*_*RPE*_ = 1.3. The final mesh configuration achieves a cup shape; **b-b’**, strain pattern resulting from an in-plane isotropic gradient of cell area increase at both layers, here *R* = 75 and *a* = 40. The final configuration exhibits the overall cup shape, but appears to acquire a shape which is a mix of the cups forming in both directions. This can be explained with lack of symmetry breaking in this pattern; **c-c’**, strain pattern resulting from an in-plane isotropic gradient of cell area increase at both layers, here *λ* = 1.1. The final mesh configuration achieves a cup shape but in the wrong direction, which can also be explained by the lack of symmetry breaking in the pattern

**Extended Fig. 5.**
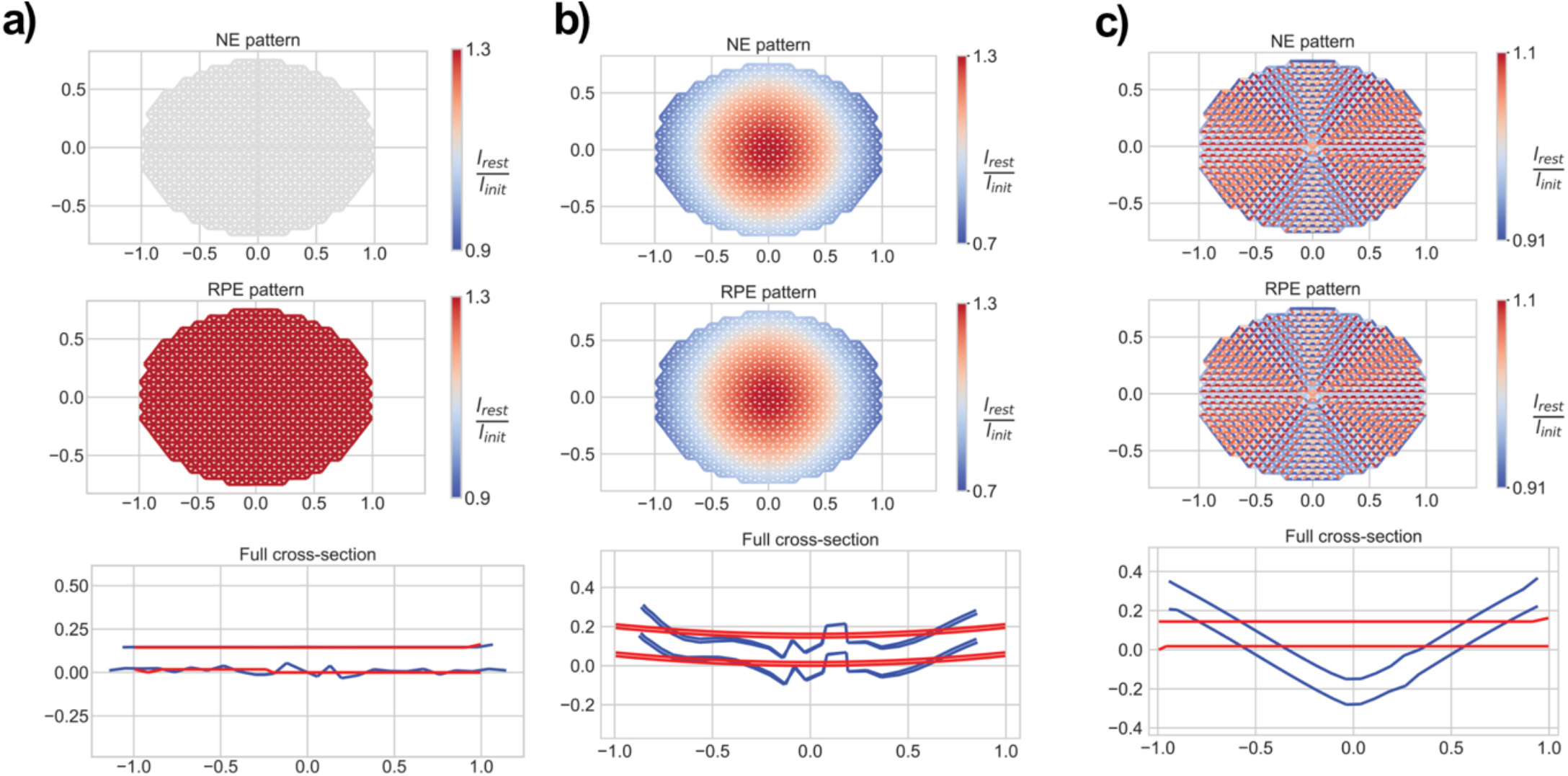
Model of ablation experiments on the simplified apical surfaces. described in the methods section. Patterns used are the same as in (**Fig. 3c-e). a**, applying the total area mismatch to the ablated mesh fails to lead to cup formation; **b-c** applying the local isotropic growth gradient and the local uniform anisotropic strain patterns to the ablated mesh both lead to cup formation.

**Extended Fig. 6.**
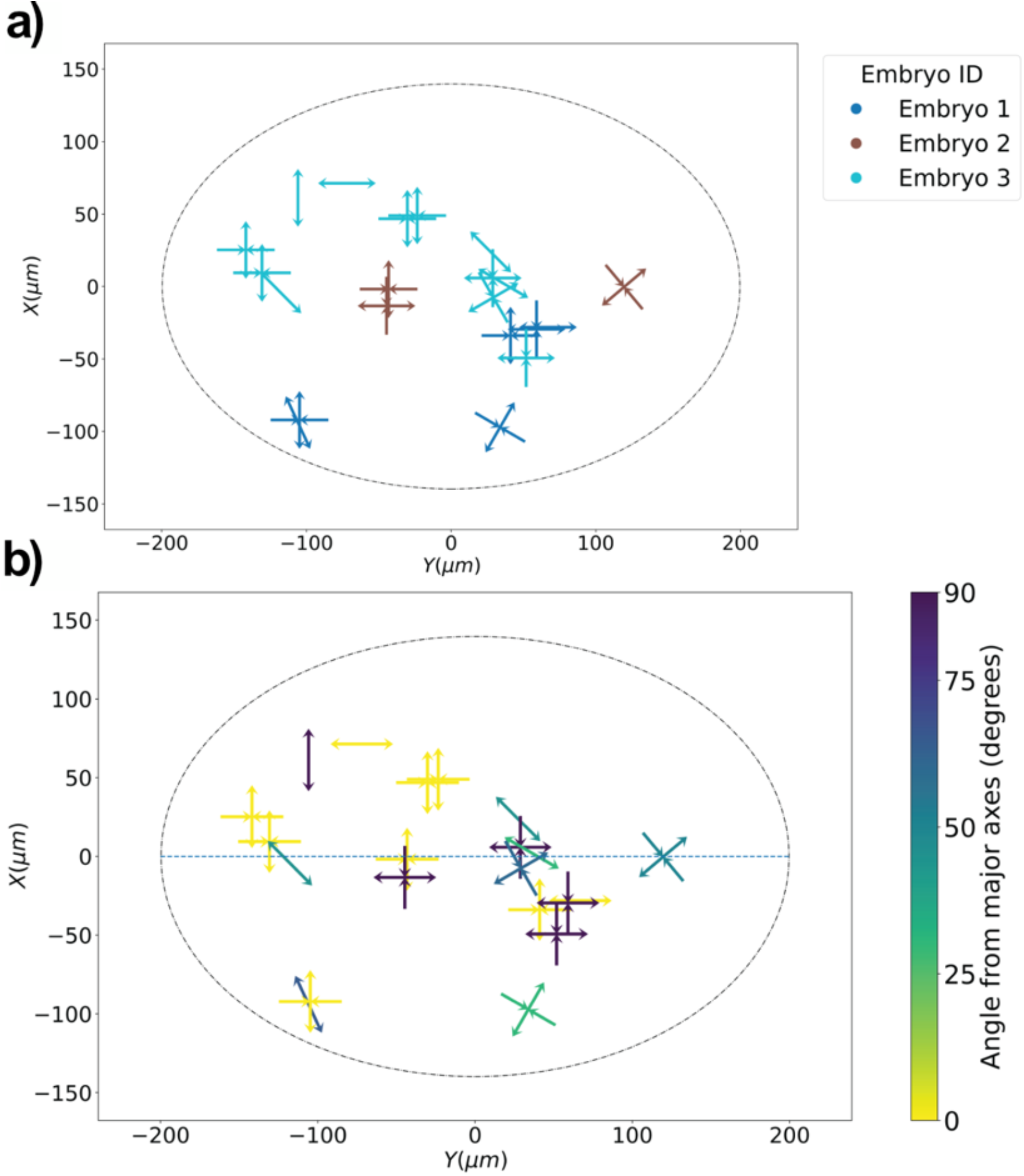
Quantification of the orientation of NE rearrangements in mosaicaly labelled embryos. **a**, collection of all events tracked in the labelled embryos by embryo ID. 2 types of events were identified – 1. rearrangements of cells marked by contracting and extending double-arrows perpendicular to each other, the contraction is aligned with the vanishing edge; 2. Appearance of new edge marked by a single extending double arrow. (n=3) **b**, quantification of angle relative to major axis of the embryo of tracked rearrangements. For neighbour exchange angle is taken from the direction of edge contraction and for edge appearance from the direction of appearance. This shows slight bias for contracting along the major axis, which coincides with an elliptical anisotropic strain pattern.

**Extended Fig. 7.**
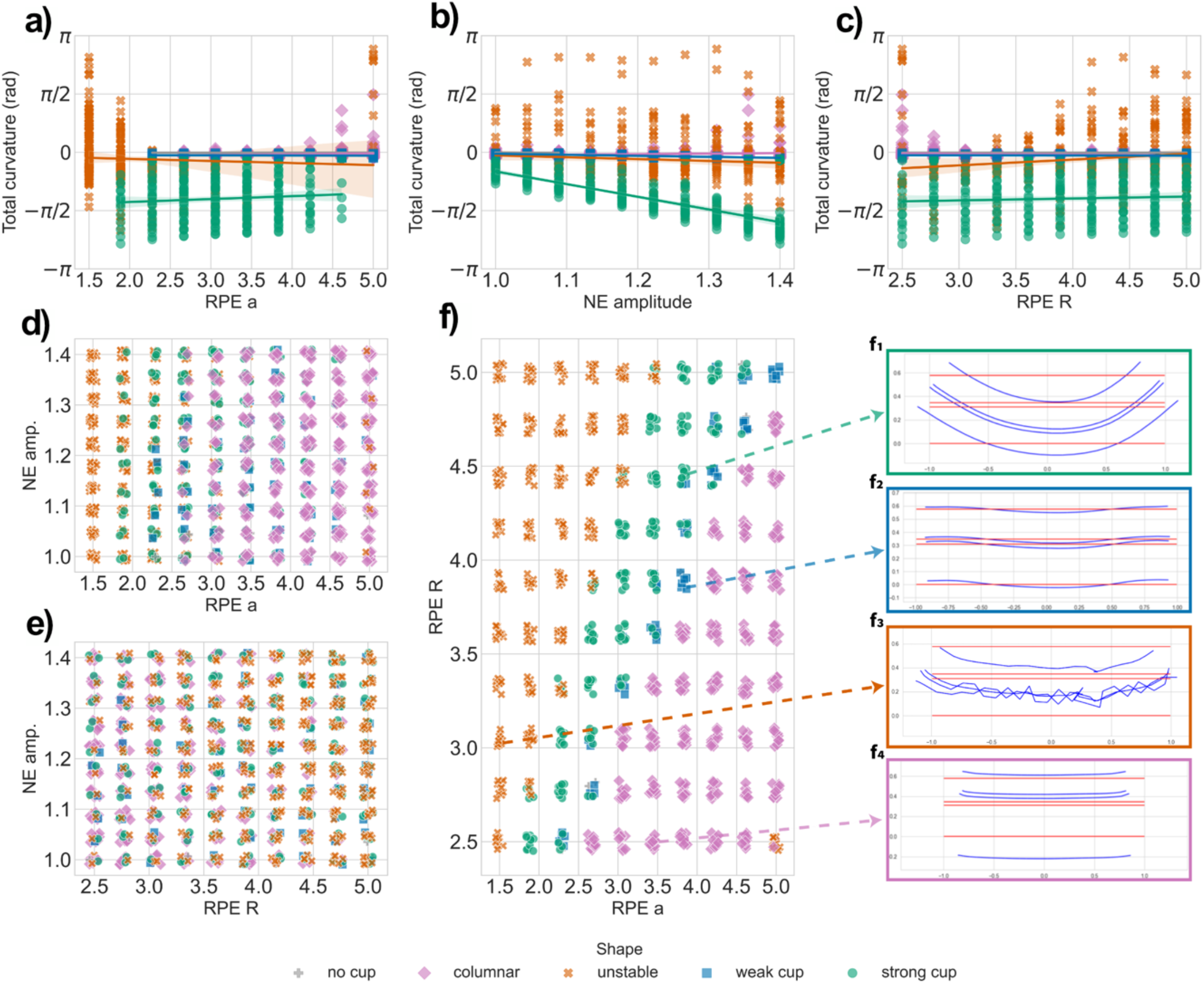
Analysis of pattern parameters sweep. quantification of x-z cross-sections of parameter sweep detailed in the methods section. Shape was determined by a set of rules on three metrics – Total basal NE cross-section curvature, standard deviation of the apical RPE cross section gradient and the minimal separation distance between the RPE apical and basal cross-sections. The shape was determined to be columnar if the minimal separation was larger than the initial separation between the layers (> 0.32*μm*) and the gradient std was low (< 0.6); Otherwise, shape was numerically unstable if the gradient std was high (> 0.25); Lastly, for shapes with low gradient std (< 0.25) shape was determined by the total curvature – flat if total curvature was > −1^°^, else weak cup if total curvature was > −10^°^and strong cup if angle was < −10^°^. **a-c**, jittered scatter plot of each of the total curvature against each of the strain patterns’ parameters coloured by shape label. No single parameter is able to distinct between the different shapes. For strong cups, NE parameter controls the amount of curvature; **d-f** phase plots of pairs of strain patterns’ parameters shows that well defined zones of shape arise in the RPE pattern phase space (**f**) – a clear boundary arises between columnar tissues and strong cup transitionins through flat and weak cups. Columnar tissues are a result of RPE pattern that has a mean local contraction instead of growth. This shows a broad region of parameters that results in a cup forming; **f1-f4** examples of the different shapes.

**Extended Fig. 8.**
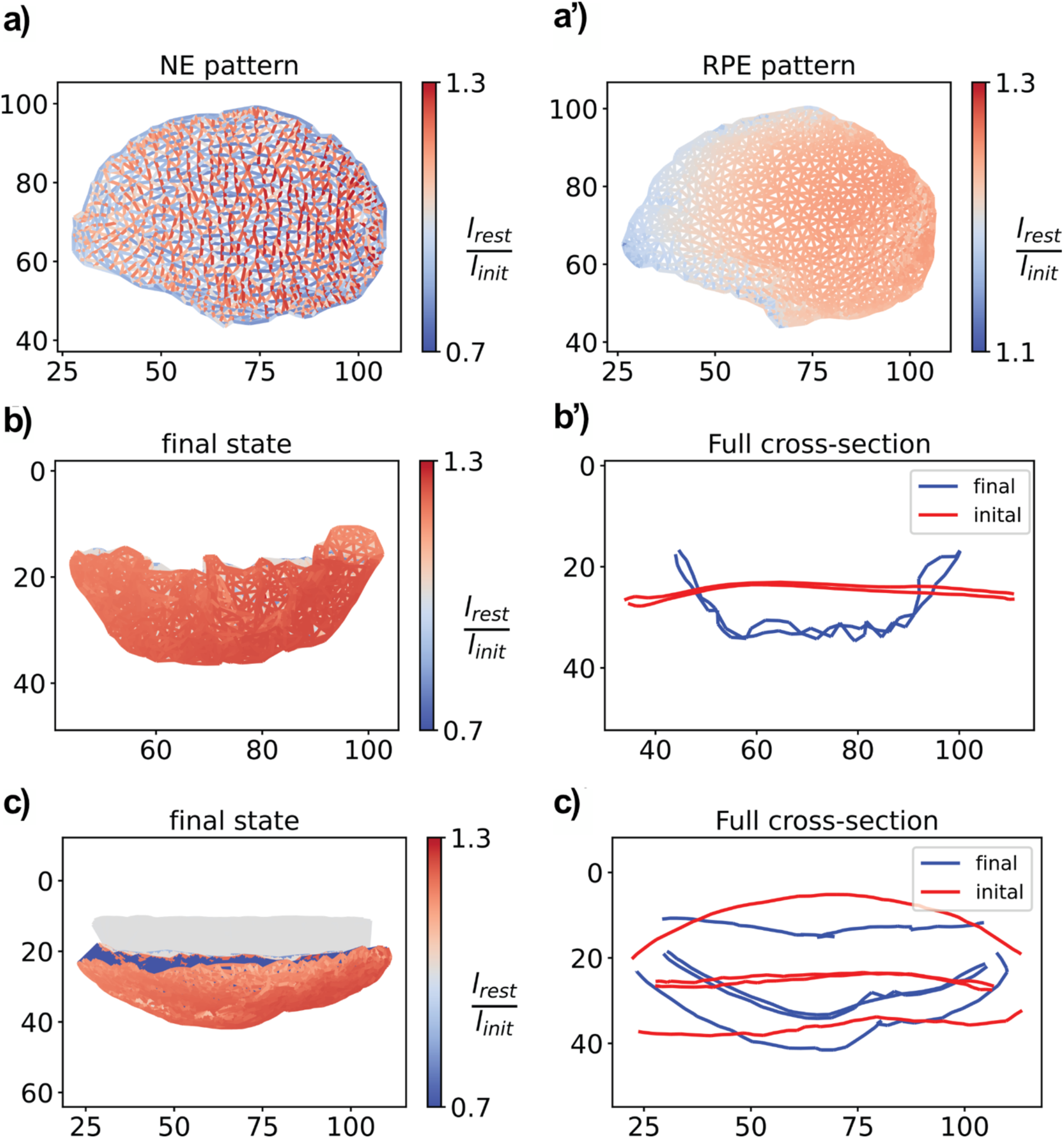
Simulation results of inferred strain pattern on segmented meshes. **a-a’**, the inferred pattern applied to the NE and RPE apical sides of the segmented meshes from the image analysis pipeline with NE *A* = 1.3 and RPE *R* = 5, *a* = 4; **b-b’**, x-z view of simulation results of applying the pattern to the isolated apical surfaces mesh, the apical sides develop into a cup in the right orientation (z-axis flipped to maintain orientation convention as RPE is above in the extracted mesh); **c-c’**, x-z view of simulation results of applying the pattern to the full segmented mesh, the apical sides develop into a cup in the right orientation causing the basal side to invaginate (z-axis flipped to maintain orientation convention as RPE is above in the extracted mesh). An inversion of the initial NE curvature occurs similar to the experimental system;

**Extended Fig. 9.**
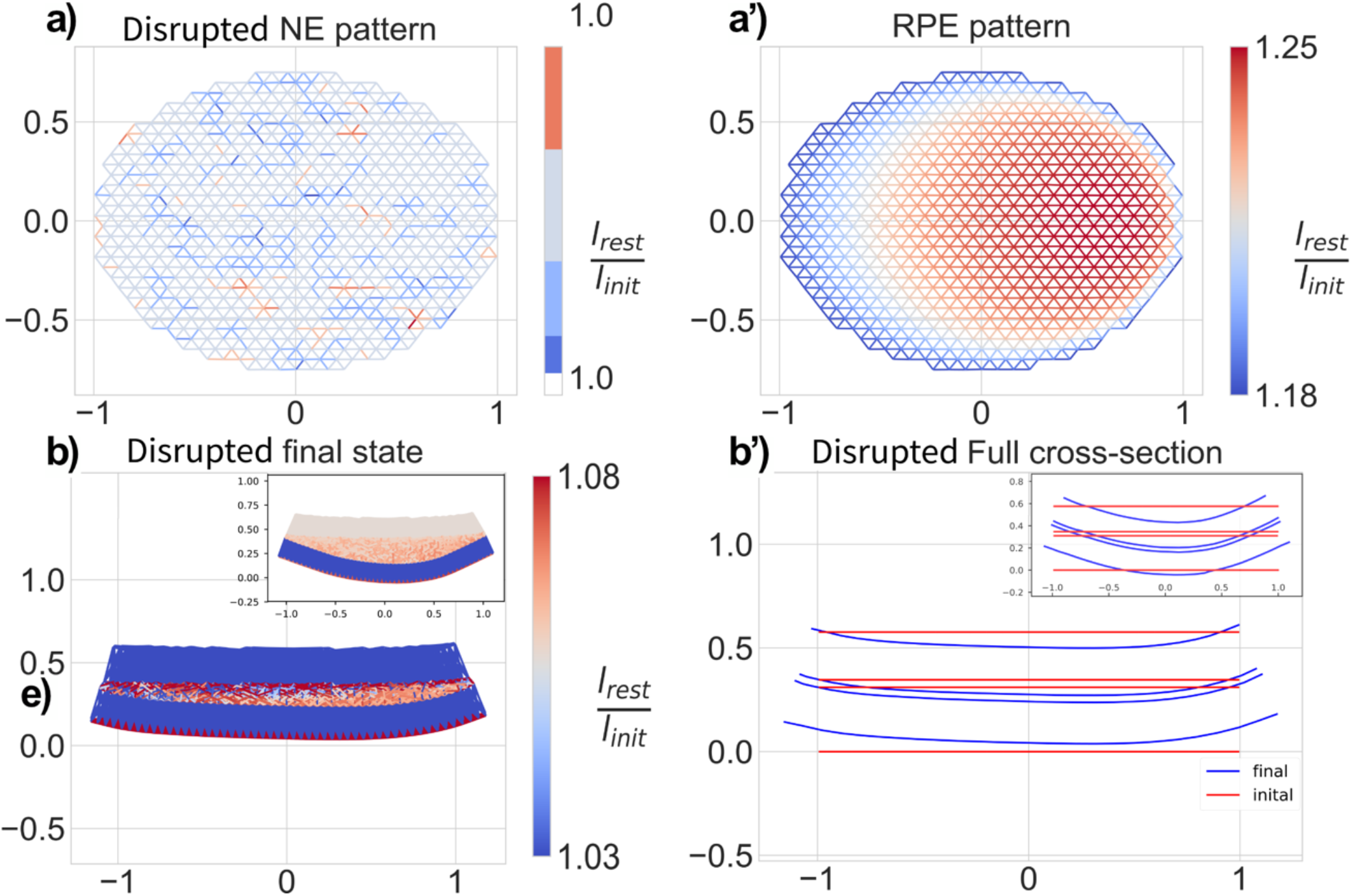
Simulation of NE pattern disruption. **a-a’**, The disrupted pattern used in the simulation: RPE pattern is identical to **(Fig 5a)** while the NE apical springs remain stress-free; **b-b’**, x-z views of the resulting shape clearly shows a reduction of final cup curvature compared to the non-perturbed case (shown in the insets).

